# Rapid ionic current phenotyping (RICP) identifies mechanistic underpinnings of iPSC-CM AP heterogeneity

**DOI:** 10.1101/2023.08.16.553521

**Authors:** Alexander P. Clark, Siyu Wei, Kristin Fullerton, Trine Krogh-Madsen, David J. Christini

**Affiliations:** Department of Biomedical Engineering, Cornell University, Ithaca, New York, USA; Department of Physiology and Pharmacology, SUNY Downstate Health Sciences University, Brooklyn, New York, USA; Department of Physiology & Biophysics, Weill Cornell Medicine, New York, New York, USA; Institute for Computational Biomedicine, Weill Cornell Medicine, New York, New York, USA

**Keywords:** Induced pluripotent stem cells, Patch clamp, Arrhythmias, Ion channels, Computer simulation

## Abstract

As a renewable, easily accessible, human-derived *in vitro* model, human induced pluripotent stem cell derived cardiomyocytes (iPSC-CMs) are a promising tool for studying arrhythmia-related factors, including cardiotoxicity and congenital proarrhythmia risks. An oft-mentioned limitation of iPSC-CMs is the abundant cell-to-cell variability in recordings of their electrical activity. Here, we develop a new method, rapid ionic current phenotyping (RICP), that utilizes a short (10 s) voltage clamp protocol to quantify cell-to-cell heterogeneity in key ionic currents. We correlate these ionic current dynamics to action potential recordings from the same cells and produce mechanistic insights into cellular heterogeneity. We present evidence that the L-type calcium current is the main determinant of upstroke velocity, rapid delayed rectifier K^+^ current is the main determinant of the maximal diastolic potential, and an outward current in the excitable range of slow delayed rectifier K^+^ is the main determinant of action potential duration. We measure an unidentified outward current in several cells at 6 mV that is not recapitulated by iPSC-CM mathematical models but contributes to determining action potential duration. In this way, our study both quantifies cell-to-cell variability in membrane potential and ionic currents, and demonstrates how the ionic current variability gives rise to action potential heterogeneity. Based on these results, we argue that iPSC-CM heterogeneity should not be viewed simply as a problem to be solved but as a model system to understand the mechanistic underpinnings of cellular variability.

**New & Noteworthy:** We present rapid ionic current phenotyping (RICP), a current quantification approach based on an optimized voltage clamp protocol. The method captures a rich snapshot of ionic currents that provides quantitative information about multiple currents (e.g., I_CaL_, I_Kr_) in the same cell. The protocol helped to identify key ionic determinants of cellular action potential heterogeneity in iPSC-CMs. This included unexpected results, such as the critical role of I_Kr_ in establishing the maximum diastolic potential.

## 1 Introduction

Human induced pluripotent stem cell derived cardiomyocytes (iPSC-CMs) are a promising model to study congenital (Terrenoire et al., 2013; Han et al., 2014) and acquired (Mathur et al., 2015) cardiac arrhythmias. iPSC-CMs can be derived from patient cells taken through a minimally invasive blood draw or skin biopsy. The resultant cells contain the genetic information of donors, and offer the potential to study patient-specific phenotypes in the lab (Liang et al., 2016).

The depth of insights from electrophysiological studies with these cells, however, is limited by their immature phenotype and cell-to-cell variability (Goversen et al., 2018b). Even genetically identical iPSC-CMs derived from the same donor display significant heterogeneity (Ma et al., 2011; Doss et al., 2012; Quach et al., 2018; Kernik et al., 2019; Akwaboah et al., 2021). Such shortcomings make it difficult to glean meaningful physiological information from patient-specific iPSC-CMs (Blinova et al., 2019), and have led to inconsistent results in multisite drug cardiotoxicity screening studies (Blinova et al., 2018). Developing an understanding of iPSC-CM heterogeneity is an essential step to improve the utility of these cells as a tool for use in precision medicine.

In recent years, extensive intercellular and intersubject variations in ion-channel expression patterns have been documented and motivated the development of cell-specific and heterogeneous populations of *in silico* models (Sarkar et al., 2012; Britton et al., 2013; Krogh-Madsen et al., 2016; Ni et al., 2018; Whittaker et al., 2020; Grandi et al., 2023; Lachaud et al., 2022; Kernik et al., 2019). Such modeling has identified electrophysiological features that increase the risk of proarrhythmic events (Passini et al., 2017; Gong and Sobie, 2018; Ni et al., 2018).

While well-documented, here we argue that iPSC-CM electrophysiological heterogeneity is a pervasive and understudied issue. We use a combination of *in vitro* and *in silico* approaches and present a new method called rapid ionic current phenotyping (RICP) that makes it possible to correlate cell-specific ionic currents to AP morphology. The approach uses a recently published voltage clamp (VC) protocol from our lab (Clark et al., 2022) — the 10 s protocol provides insight into the presence and relative size of several key cardiac ionic currents. Using this data, we identify currents that influence AP morphology and explain both cell-to-cell variability and outliers in a population of iPSC-CMs.

We also use the short, 10 s, VC protocol data to draw the following conclusions about the cells used in this study:

- L-type calcium current (I_CaL_) drives upstroke in iPSC-CMs with a depolarized AP morphology.
- Rapid delayed rectifier K^+^ current (I_Kr_) plays a role in establishing the maximal diastolic potential.
- Seal-leak current contaminates VC recordings and contributes to a depolarized maximal diastolic potential.
- Large positive currents present at potentials >40 mV correlate with a shortened AP duration.
- Several cells contain a strong outward current at 6 mV that is not present in computational AP models and correlates with AP duration.

## 2 Methods

### 2.1 iPSC-CM cell culture and electrophysiological setup

The *in vitro* data was previously published (Clark et al., 2022).

Frozen stocks of human iPSC-CMs were purchased from the Stanford Cardiovascular Institute Biobank. These cells were derived from an African-American female donor in a process approved by Stanford University Human Subjects Research Institutional Review Board. The use of cells from one donor allowed us to develop a method that aims to explain heterogeneity within a single individual, rather than considering effects at the population level.

Cells were prepared for electrophysiological experiments following the steps described in Clark et al. (2022). Briefly, cells were thawed and cultured as a monolayer in one well of a 6-well plate precoated with 1% Matrigel. Cells were cultured with RPMI media (Fisher/Corning 10-040-CM) containing 5% FBS and 2% B27 and kept in an incubator at 37 °C, 5% CO_2_, and 85% humidity. After 48 hours, cells were lifted with 1 mL Accutase, diluted to 100, 000 cells*/*mL, and replated on 124 sterile 8 mm coverslips precoated with 1% Matrigel. Cells were cultured with RPMI media that was swapped every 48 hours. Cells were patched between days 5 and 15 after thaw. Voltage clamp and current clamp recordings were acquired from 40 cells using the perforated patch technique (see Clark et al. (2022) for details). We excluded one cell from the analyses in this study because it had spontaneous alternans with inconsistent AP features. All cells had a pre-rupture seal of >300 MΩ.

### 2.2 Voltage clamp protocol

We have previously developed a voltage clamp protocol consisting of multiple short segments, each designed to isolate one key ionic current (Clark et al., 2022). The protocol was designed using optimization techniques and a mathematical model of iPSC-CMs (Kernik et al., 2019) to maximize, one at a time, the contribution to total current by each of seven key currents: I_Kr_, I_CaL_, sodium current (I_Na_), transient outward K^+^ current (I_to_), inward rectifier K^+^ current (I_K1_), funny current (I_f_), and slow delayed rectifier K^+^ current (I_Ks_).

During our analysis, we found that the current measured 100 ms after a depolarizing step to 6 mV (I_6mV_) was substantially different in many cells from that predicted by the mathematical model and we therefore included I_6mV_ as an 8th current measure.

For each current-isolating segment, we quantified the recorded total current I_out_ using either the minimum or the average over a 2 ms span centered at the following values (Figure 3C): I_6mV_ (600 ms, average), I_Kr_ (1262 ms, average), I_CaL_ (1986 ms, minimum), I_Na_ (2760 ms, minimum), I_to_ (3641 ms, average), I_K1_ (4300 ms, average), I_f_ (5840 ms, average), and I_Ks_ (9040 ms, average).

**Figure 1:**
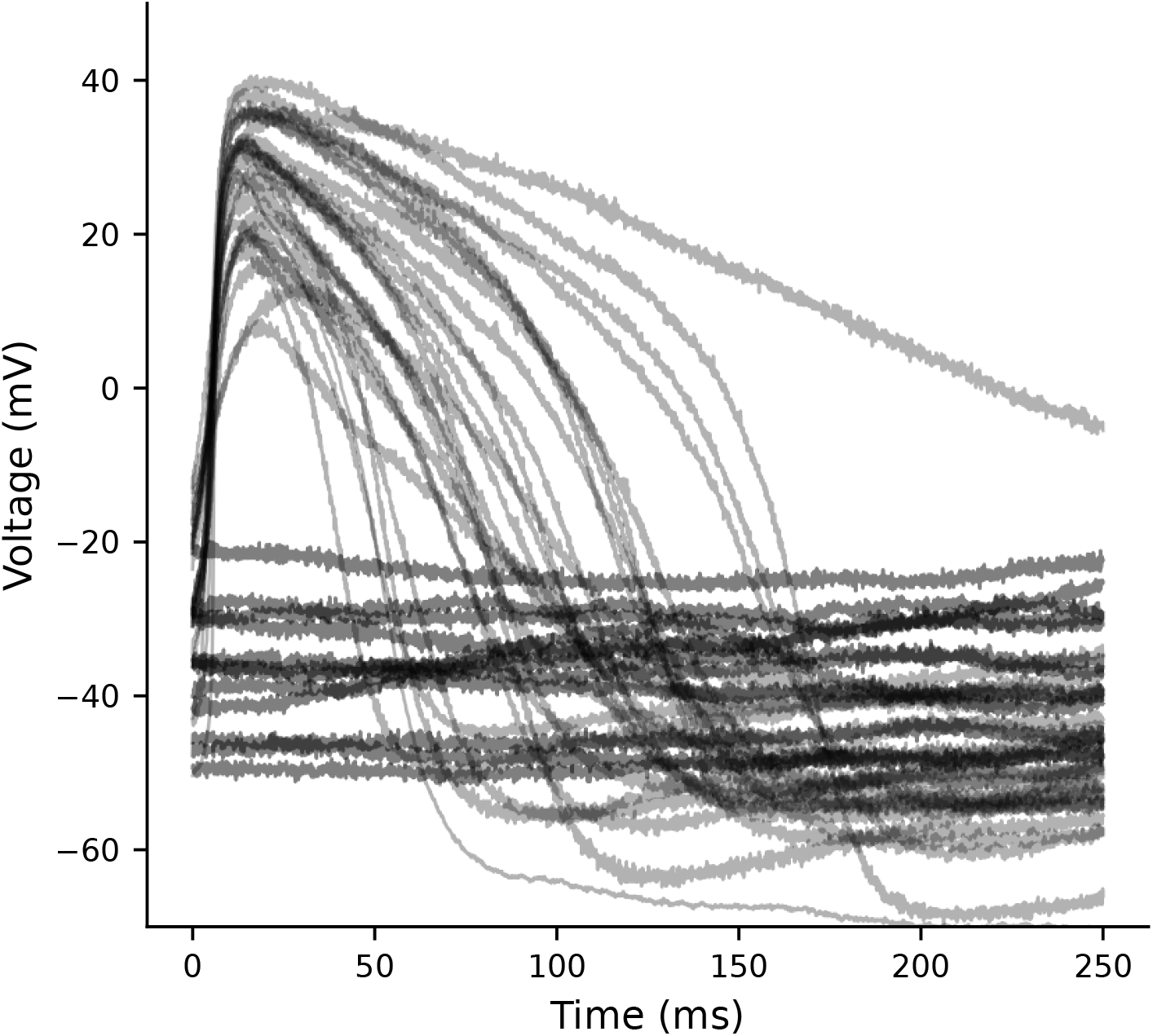
Current clamp recordings illustrate baseline heterogeneity. Spontaneously beating (n=27) and quiescent depolarized cells (n=12) from current clamp recordings of iPSC-CMs. If reported as *mean*± *SEM*, the variation in MP (−52 ± 1.6 mV), APD_90_ (132 ± 14 ms), and dV/dt_max_ (10.4 ± 1.2 V/s) appears quite small.

**Figure 2:**
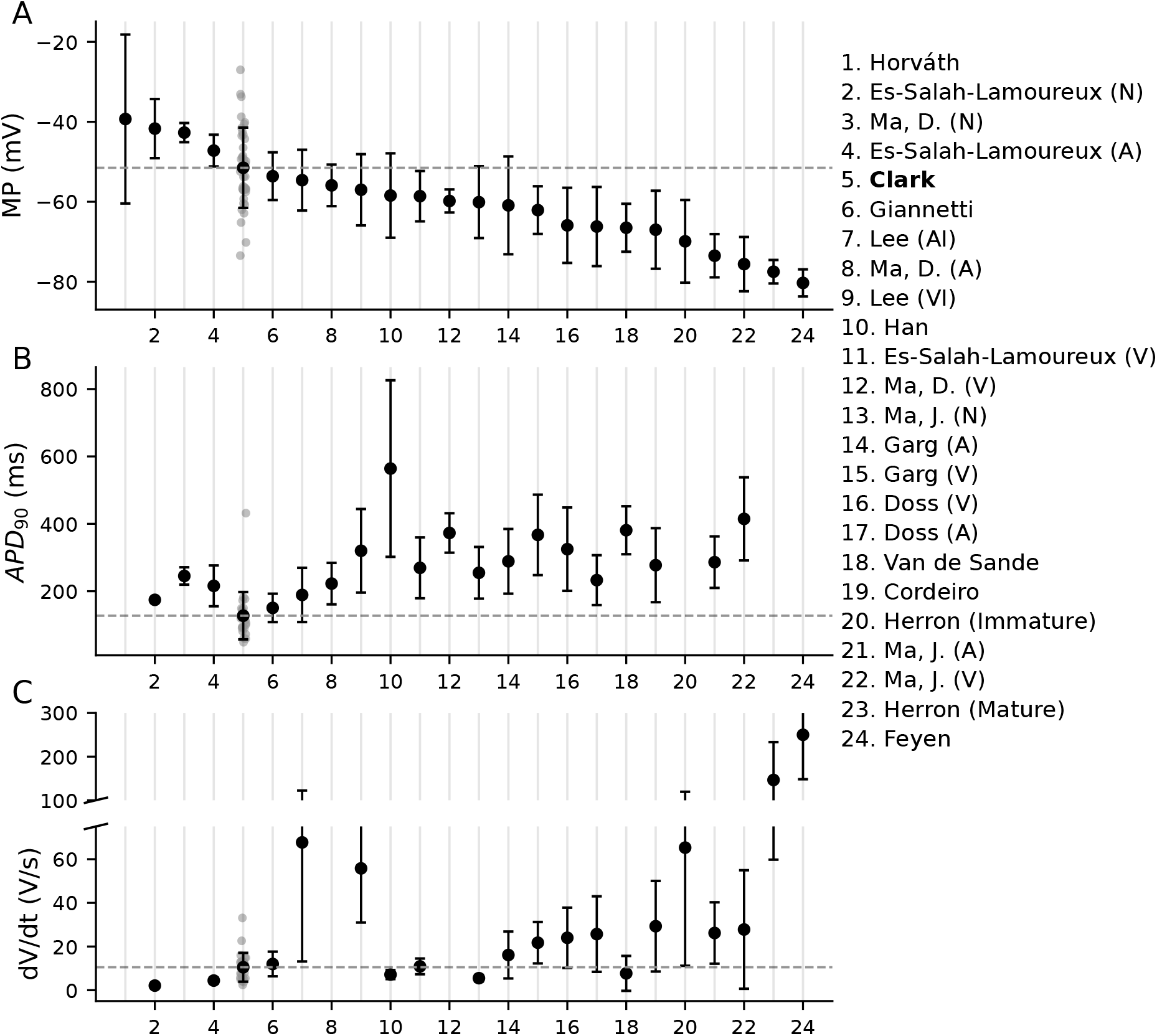
Variations in AP features from 24 independent datasets. The numbered studies on the right are independent datasets, named according to the first author (see references for full bibliographic information), and the numbers correspond to their position on the x-axis. Studies labeled with (N), (A), or (V) used AP features to sort cells into nodal, atrial, or ventricular groupings. The datasets from Lee et al. (2017) used differentiation protocols to induce either atrial-like (AI) or ventricular-like (VI) phenotypes. The datasets in this figure, including our own (position 5), are sorted in order from most to least depolarized MP.

**Figure 3:**
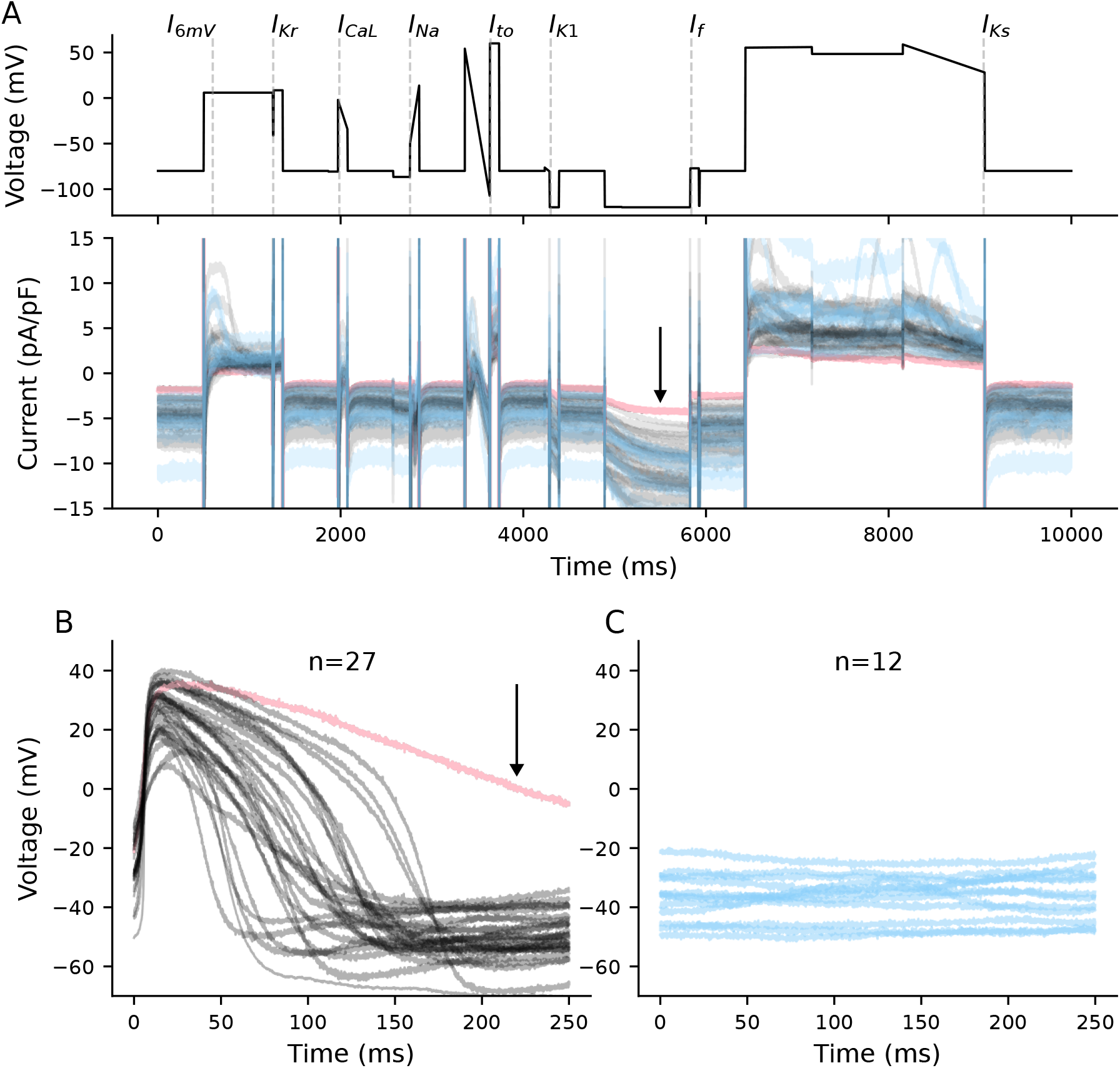
VC variations and membrane potential in 39 cells. **A**, The optimized voltage clamp protocol (top), with lines overlaid that indicate time points designed to isolate each current. Responses to the voltage clamp protocol (bottom) for spontaneous (black), quiescent (blue), and the outlier (pink). **B**, APs from spontaneously beating cells (n=27). The pink AP is an outlier with an APD_90_ >2x longer than the rest of the population. **C**, Voltage recordings from quiescent and depolarized cells (n=12). The voltage recordings in **B** and **C** are the same as those shown in Figure 1 but are here separated for clarity.

### 2.3 AP feature calculations

The transmembrane potential of each cell was recorded for 10 s. Of the 39 cells, 12 were not spontaneously beating. We computed a minimal potential (MP) for these non-spontaneous cells.

For cells that were spontaneously beating, we computed their MP (in this case, the minimum voltage during the AP), action potential duration at 90% repolarization (APD_90_), cycle length (CL), and maximal upstroke velocity (dV/dt_max_). The average of each feature was calculated for all cells that produced more than one AP during the 10 s recording.

### 2.4 iPSC-CM mathematical models

For comparison and to guide the analysis of our experimental data, we used two different mathematical models of iPSC-CM electrophysiology: the Paci et al. model (Paci et al., 2013) and the Kernik et al. model (Kernik et al., 2019). We set the cell capacitance (C_m_) of these models to 45 pF, which is centrally located in the range (18-98 pF) of the capacitances for cells used in this study.

To avoid long transients and to better simulate our perforated patch experimental setup, we fixed intracellular sodium and potassium concentrations ([Na^+^]_i_ and [K^+^]_i_) to their baseline steady state values (taken after 1000 s of spontaneous or paced current clamp simulation).

Because the leak through the imperfect pipette-membrane seal during single-cell patch-clamp experiments can substantially impact the electrophysiological recordings in these cells, we included a linear leak current (I_leak_) in the mathematical models (Clark et al., 2023). We used a baseline value of 2 GΩ for the seal resistance.

In addition to the seal-leak current, for voltage clamp simulations, we included explicit modeling of the following experimental artifacts: liquid junction potential offset (-2.8 mV), access resistance (20 MΩ), and series resistance (R_s_) compensation (70%), including supercharging (Lei et al., 2020; Lei, 2020).

### 2.5 Population of models and sensitivity analysis

A population of 500 individual models was generated from both the Paci and Kernik models with experimental artifact equations by randomly sampling conductances of I_Na_, I_CaL_, I_Kr_, I_Ks_, I_to_, I_K1_, I_f_, I_leak_, sodium calcium exchange current (I_NaCa_), sodium potassium pump current (I_NaK_), sodium background current (I_bNa_), calcium background current (I_bCa_), as well as C_m_ and R_s_ between 0.25 and 4x their baseline values.

We used a Spearman correlation to determine the sensitivity of the current-isolating time points to these parameters.

### 2.6 Linear regression

A linear least-squares regression was used to compare ionic currents and AP features for both the *in vitro* and the *in silico* data. A Spearman correlation coefficient and p-value were calculated for these data.

### 2.7 Software and simulations

Simulations were performed in Myokit v1.33.7 (Clerx et al., 2016). Additional analysis was done in Python using NumPy v1.21.6 and SciPy v1.7.3 (Virtanen et al., 2020).

All data, code and models can be accessed from GitHub (https://github.com/Christini-Lab/ap-vc-correlations.git).

## 3 Results

### 3.1 iPSC-CMs are heterogeneous

Of the 39 cells used in this study, 27 were spontaneously beating, and 12 were quiescent and depolarized (Figure 1). There is substantial cell-to-cell heterogeneity. Reporting the variation as the standard deviation (SD), we calculate a MP of −52 ± 10 mV, action potential duration at 90% repolarization (APD_90_) of 127 ± 70 ms, and maximum upstroke velocity (dV/dt_max_) of 10.5 ± 6.6 V/s. Within the subset of spontaneously beating cells, there is additional heterogeneity in that some have a predictable and consistent CL, while the CL of others is highly variable (Figure S1)

Large cell-to-cell variations in iPSC-CM AP data is common in the literature. Figure 2 shows the mean and SD error bars for MP, APD_90_, and dV/dt_max_ of 24 independent datasets taken from 14 studies. We selected these three AP features because they were the most consistently reported in manuscripts and have straightforward operational definitions. These data illustrate the large inter- and intralab heterogeneity present in iPSC-CM studies.

### 3.2 Rapid ionic current phenotyping provides insight into AP outliers

As detailed in the Methods, we recently published a voltage clamp protocol (Clark et al., 2022) that was designed to isolate each of the following seven key ionic currents: I_Kr_, I_CaL_, I_Na_, I_to_, I_K1_, I_f_, and I_Ks_. The protocol works by stepping to voltages designed to maximize the contribution of each current at different time points. Because of its short duration (10 s) and design to target multiple currents, we are calling this approach rapid ionic current phenotyping (RICP).

Figure 3A shows the voltage clamp protocol (top) and heterogeneity in ionic current responses (bottom) from the 39 cells included in this study. On the voltage clamp protocol, we have overlaid dashed lines to highlight current-isolating time points that we use to study ionic current dynamics. In addition to the seven designed current-isolating segments, we included another time point measure (I_6mV_, the current recorded 100 ms after a depolarizing step to 6 mV), as we found that it provides insight into dynamics that are not present during other portions of the voltage clamp protocol.

Spontaneously beating cells are plotted in black and quiescent cells are in blue (Figure 3A-C). The pink spontaneous cell is an outlier, with an APD_90_ value >2x longer than the rest of the population.

Figure 4 shows the distribution of recorded total current values at each of the eight current-isolating time points. Similar to the AP data, we see a large variation in measurements during each of these current-isolating segments. Compared to the rest of the population, the outlier cell appears to have the smallest total current during the regions of the protocol designed to isolate I_K1_, I_f_, and I_Ks_. All three of these currents conduct potassium ions that often contribute to repolarizing cardiomyocytes. This provides a possible explanation for how a cell with current measures at the tail ends of the distributions, can become an outlier in terms of an AP metric (e.g., APD), and shows how one may mechanistically explain outlier voltage dynamics in terms of the underlying ionic currents.

**Figure 4:**
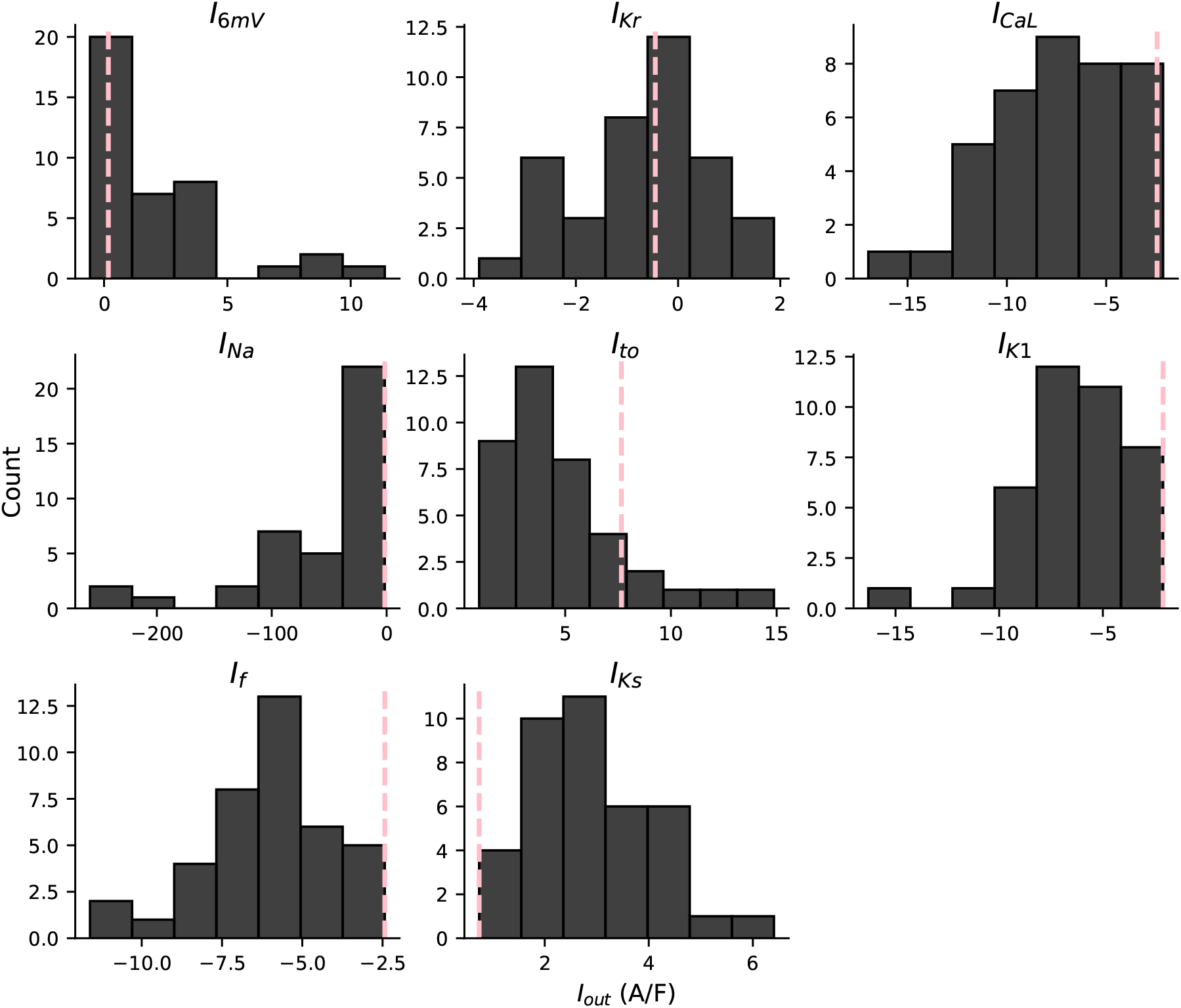
I_out_ distribution at the eight current-isolating time points. Histograms display the distribution of recorded current values at each of the eight time points. The pink dashed line shows I_out_ values for the outlier cell.

### 3.3 RICP identifies I_CaL_ as driver of upstroke in depolarized cells

Figure 5A displays the relationship between the total current during each current-isolating segment and the dV/dt_max_ for all cells. None of the eight current-isolating segments correlate with dV/dt_max_. Figure 4 shows the likely presence of I_Na_ in many of these cells — we draw this conclusion because I_Na_ is the only current expected to generate an I_out_ of *<* −40 A/F within 2 ms after the I_Na_ voltage step. However, the upstroke velocity is relatively small in most cells (Figure 2C) and does not correlate with the current measured I_Na_-isolating segment. The small upstroke velocities and the lack of correlation with I_Na_ was an unexpected finding, given that I_Na_ is present in many of these cells.

**Figure 5:**
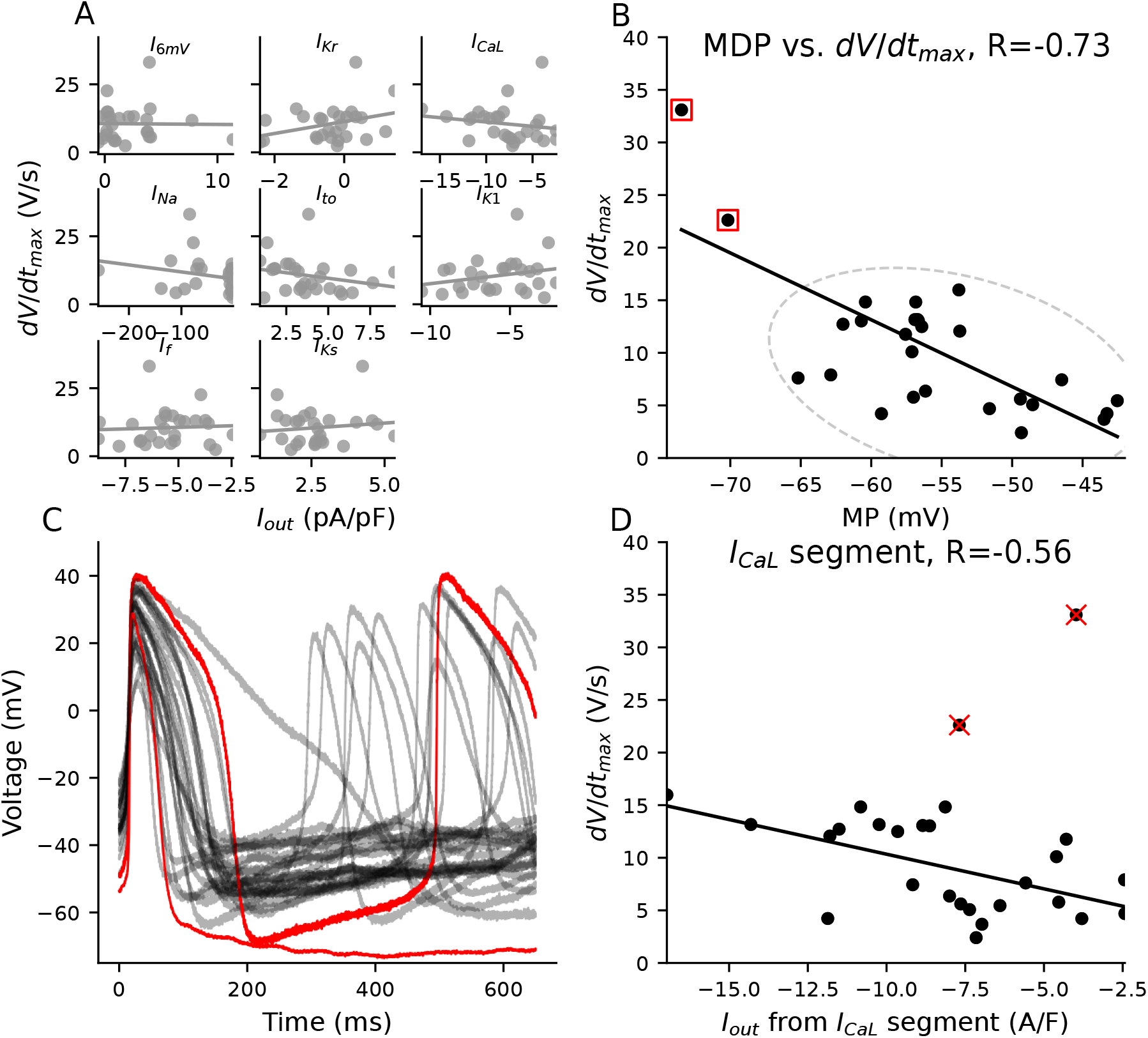
I_CaL_ helps drive upstroke in depolarized iPSC-CMs. **A**, There is no correlation between I_out_ from the eight current-isolating segments and dV/dt_max_. **B**, dV/dt_max_ decreases as MP depolarizes. The red boxes denote the two cells with the most hyperpolarized MP and largest dV/dt_max_. **C**, The two highlighted cells from panel **B** share few AP commonalities with one another other than their relatively hyperpolarized MP and large dV/dt_max_. **D**, A trend emerges between I_out_ during the I_CaL_-isolating segment and dV/dt_max_ when the two hyperpolarized cells are removed from the analysis.

While there is no significant relationship with the VC segments, dV/dt_max_ does correlate with MP (Figure 5B) — dV/dt_max_ increases as MP becomes more hyperpolarized. The two cells with MP below -70 mV (denoted with red squares) stand out as having much larger upstroke velocities than the rest of the population. I_Na_, which is responsible for the upstroke in adult ventricular and atrial cardiomyocytes, recovers from inactivation at voltages below -65 mV. As such, one well-supported hypothesis for the larger dV/dt_max_ in these two iPSC-CMs (Rook et al., 1999; Goversen et al., 2018b) is that they repolarize enough to make some sodium channels available for an I_Na_-driven upstroke. Other than having relatively hyperpolarized MP values, these two cells produce APs with few morphological similarities (Figure 5C).

Interestingly, when we exclude these cells from the regression analysis, and only consider iPSC-CMs with MP > -70 mV, a significant relationship emerges between the I_CaL_-isolating segment and dV/dt_max_ (Figure 5D).This indicates that I_CaL_ may be, at least partly, responsible for the upstroke in cells with MP>-70 mV.

### 3.4 RICP identifies I_Kr_-isolating segment as predictor of MP

Four current-isolating segments (I_Kr_, I_to_, I_K1_, and I_f_) of the VC protocol correlate with MP (Figure 6A), with the I_Kr_-isolating segment being the strongest correlate (R=0.72). We selected cells at the two MP extremes to illustrate the relationship between the I_Kr_-isolating segment and MP (Figure 6B): Cell 1 (blue) is the most hyperpolarized in the population (MP=-73 mV and APD_90_=81 ms) and Cell 2 is the most depolarized (quiescent with MP of -27 mV).

**Figure 6:**
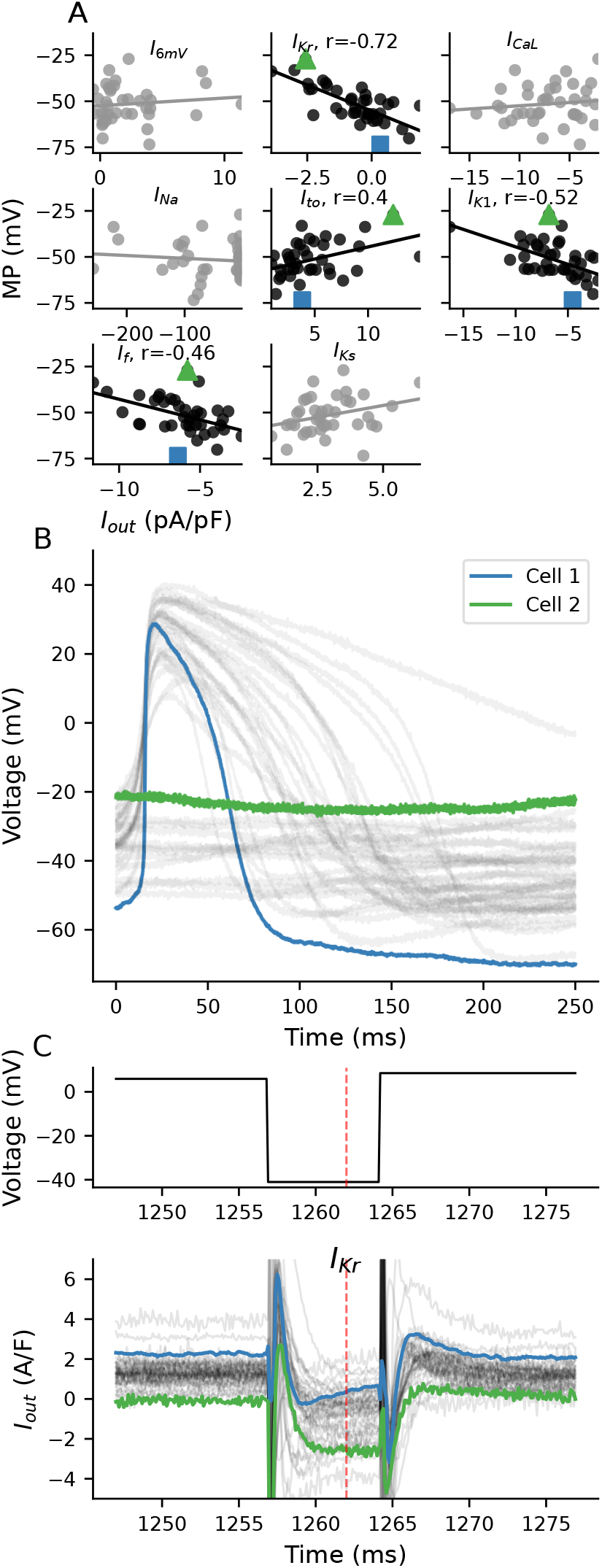
Correlations between ionic current segments and MP. **A**, I_out_ from the I_Kr_, I_to_, I_K1_, I_f_, and I_Ks_ segments correlate significantly (p*<*.05) with MP. **B**, Current clamp recordings from all cells. The iPSC-CMs with the most hyperpolarized (blue, Cell 1) and depolarized (green, Cell 2) MP values are compared. **C**, Traces from the I_Kr_-isolating segment of the VC protocol for all cells. The VC protocol (top) includes a 750 ms prestep at 6 mV, followed by a 7 ms step at -41 mV, and then a step to 9 mV. The VC protocol was designed to isolate I_Kr_ at 1262 ms (red dashed line). Cell 1 (blue) and Cell 2 (green) are highlighted.

A plot of the I_Kr_-isolating section of the VC protocol shows the voltage command (top) and current responses (bottom) for all cells in the population (Figure 6C). During this section of the protocol, cells were clamped to 6 mV for 750 ms, then stepped to -41 mV for 7 ms, and then back up to 9 mV. At the time point designed to isolate I_Kr_ (red dashed line), Cell 1 conducts a small positive total current (0.3 A/F) that increases in size, which is consistent with I_Kr_ recovering from rectification. In contrast, Cell 2 conducts a net negative current (-2.5 A/F) and does not show I_Kr_ recovery characteristics, suggesting that this cell has much less I_Kr_. These cells are examples of the population-level correlation seen between the I_Kr_-isolating segment and MP. They indicate that I_Kr_, and other currents that may be present during the step to -40 mV, likely play a critical role in establishing MP in these cells.

We used an iPSC-CM mathematical model with a linear seal-leak current (see Methods) to further study the potential role of I_Kr_ in establishing a MP. Figure 7 shows the effect of scaling I_Kr_ conductance (g_Kr_) on the MP for a model with a 2 GΩ seal resistance. While holding all other parameters constant, reducing g_Kr_ by >70% of baseline results in substantial depolarization. The 10% to 90% range of our experimental data corresponds to a roughly 70% to 85% reduction in g_Kr_ (Figure 7B). These model findings are consistent with the correlations seen between the experimental I_Kr_-isolating I_out_ measurements and MP.

**Figure 7:**
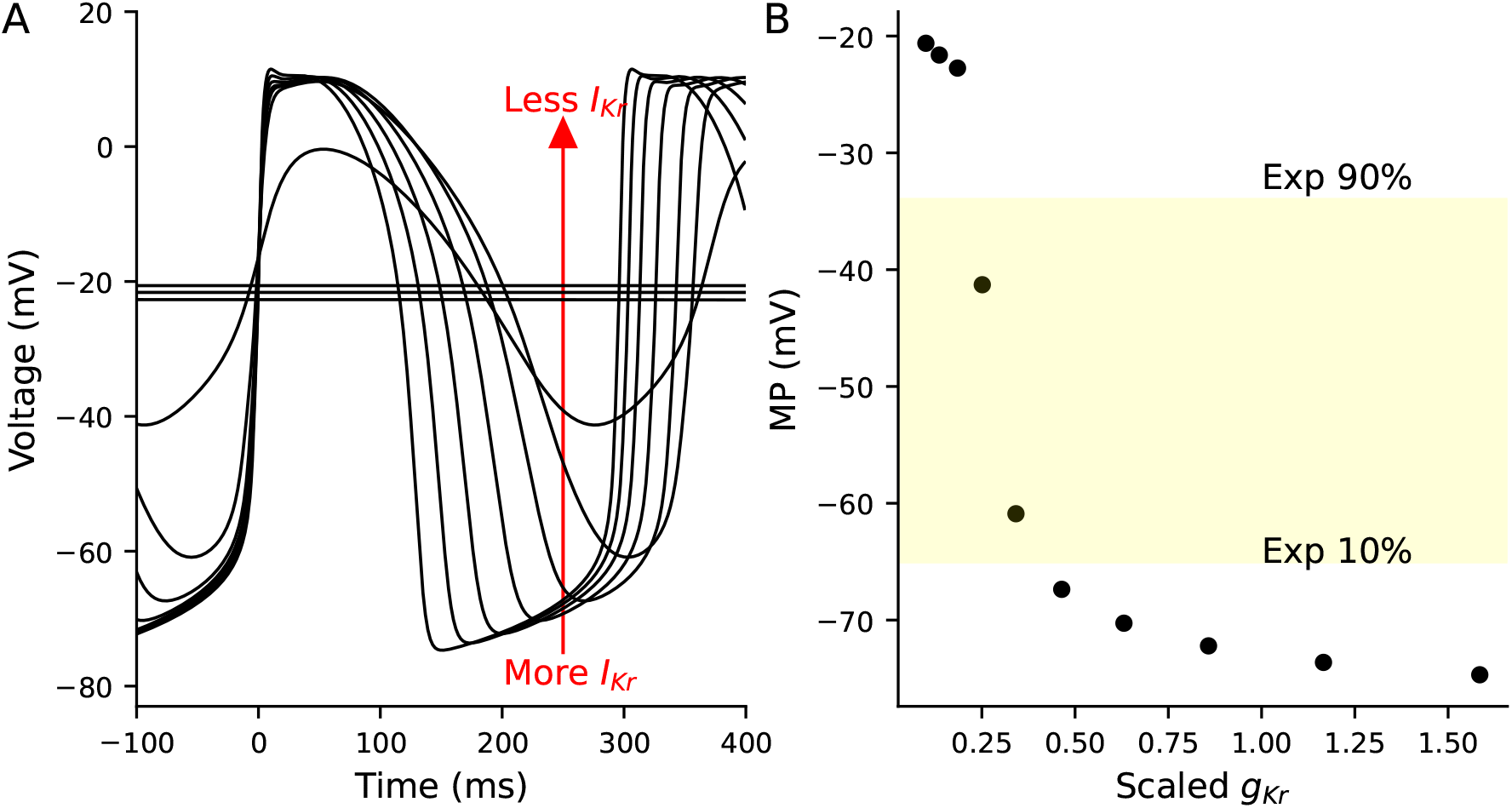
AP simulations of a Kernik-Clancy model with a 2 GΩ seal resistance and varying levels of g_Kr_. **A**, APs generated by running the model with varying levels of g_Kr_ from 0.1 to 1.6 times the baseline Kernik-Clancy value (0.218025 nS/pF). The arrow indicates that depolarized cells have less I_Kr_ (i.e., smaller g_Kr_). **B**, Relationship between g_Kr_ and MP for cells plotted in **A**. The yellow highlighted region represents the 10% to 90% range of MP values from cells in this study.

### 3.5 Seal-leak current likely contributes to the depolarized MP

In addition to the I_Kr_-isolating segment, the I_to_, I_K1_, I_f_, and I_Ks_-isolating segments also weakly correlate with MP (Figure 6A). Some of these correlations are unexpected and counter-intuitive as, e.g., I_to_ is typically inactivated during phases 3 and 4 of the AP. We hypothesized that these correlations may be influenced by a leak artifact current caused by an imperfect seal between the pipette tip and cell membrane — we recently demonstrated that this leak current, present at voltages far from zero, can substantially depolarize the MP of iPSC-CMs (Clark et al., 2023).

In an attempt to investigate the role of I_leak_ on these segments, we developed a population of Paci models that include patch-clamp experimental artifact equations (Figure 8). In this *in silico* population, the I_Kr_ segment is also the main correlate with MP, with I_to_, I_Ks_, and I_f_, being more weakly correlated with MP (figure S3).

**Figure 8:**
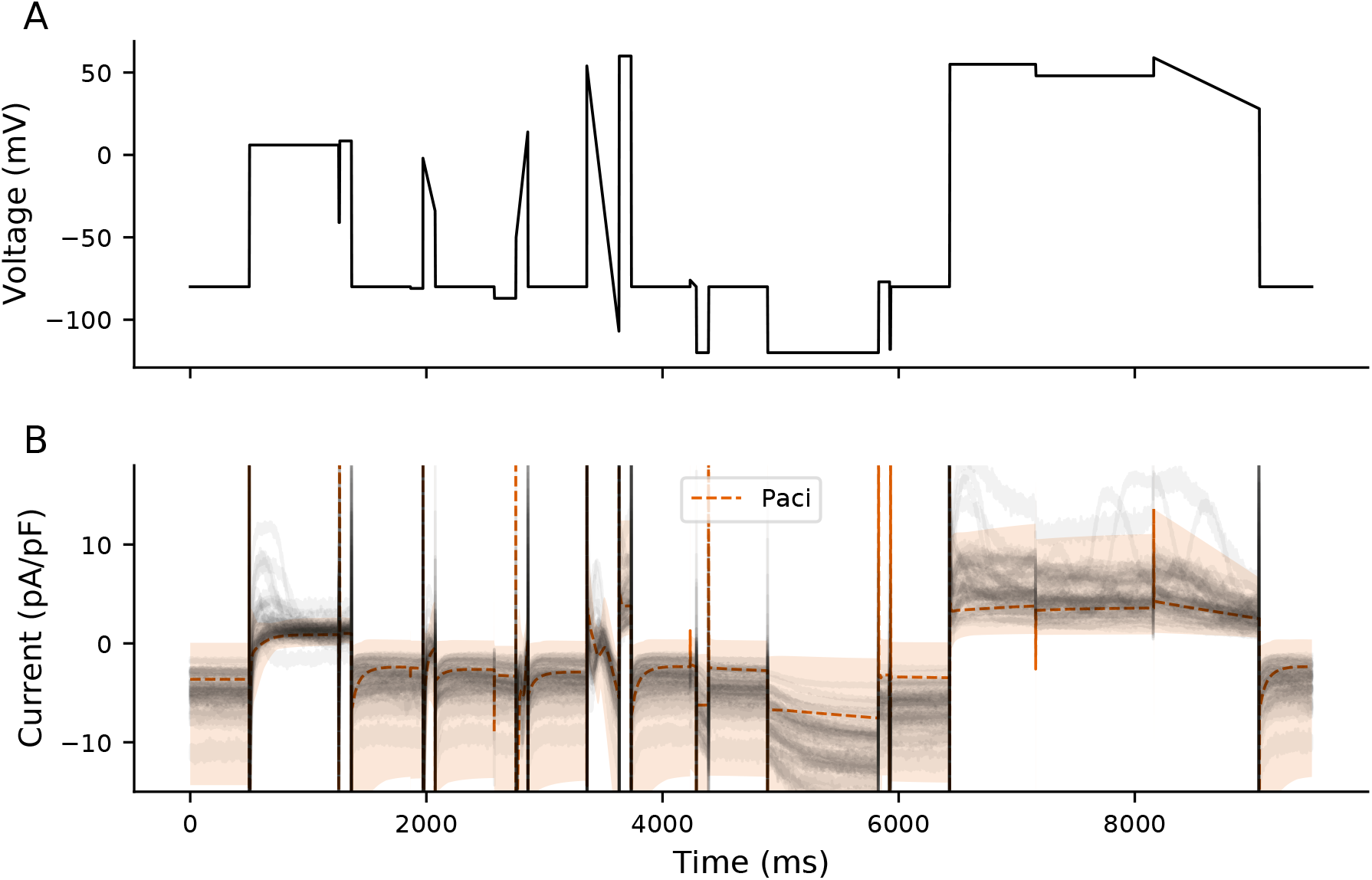
A population of Paci models compared to experimental data. 500 Paci individuals were generated by varying ionic current conductances and experimental artifact parameters (see Methods). The dashed orange line shows the average I_out_ for all models, and the shaded region is the range of values for all models. The experimental data are displayed in gray.

A sensitivity analysis of the population of models shows that the segments weakly correlating with MP in the experiments (i.e., the I_to_, I_K1_, I_f_, and I_Ks_ segments) are all sensitive to the conductance of I_leak_ (i.e., g_leak_, Figure 9). I_leak_ is modeled as a linear current with a reversal potential of zero, and so it will increase when cells are clamped to voltages that are far from 0 mV. Seeing as I_Ks_ and I_to_ are elicited by stepping to large positive voltages and I_K1_ and I_f_ to large negative voltages, this is consistent with these segments being contaminated by I_leak_. Increased g_leak_ increases the total (net outward) currents computed at positive voltages (e.g., I_Ks_ and I_to_ segments) and depolarizes the MP, consistent with the positive correlation seen between MP and the I_Ks_ and I_to_ segments in our experiments. Similarly, the negative correlations between MP and I_K1_ and I_f_ in our cells is consistent with the *in silico* results where increased g_leak_ during these hyperpolarized segments in the models contribute to the large (net inward) current and depolarized MP values.

**Figure 9:**
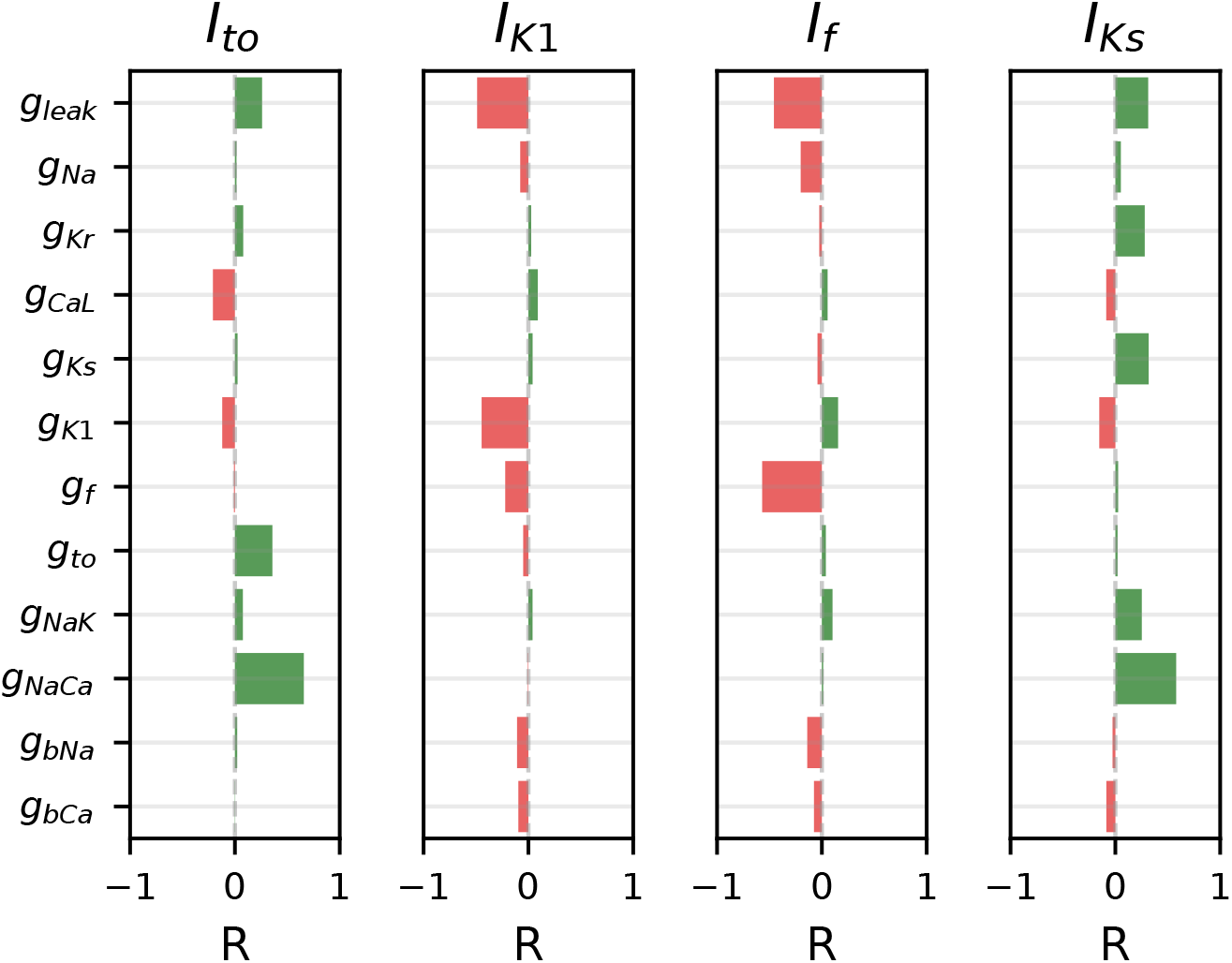
I_to_, I_K1_, I_f_, and I_Ks_-isolating segments are sensitive to seal-leak current. Correlations calculated between model parameter values (primarily conductances; e.g., g_Kr_) and I_out_ during each of these four segments. Each current is sensitive to its own conductance, with outward currents (I_to_ and I_Ks_) having a positive correlation and inward currents (I_K1_ and I_f_) showing as a negative correlation to their own conductance. Each current is also sensitive to g_leak_. Both I_to_ and I_Ks_ have a surprisingly strong sensitivity to g_NaCa_.

### 3.6 RICP identifies strong outward currents as drivers of APD_90_

Four segments (I_6mV_, I_K1_, I_f_, and I_Ks_) of the VC protocol correlate with APD_90_ (Figure 10A). Three of the segments (I_K1_, I_f_, I_Ks_) were designed to isolate potassium-conducting currents, each of which are clamped to voltages far from 0 mV. The I_6mV_ segment was added to quantify observed current dynamics different from those produced by the mathematical models and which we hypothesized could be explanatory of AP morphological differences.

**Figure 10:**
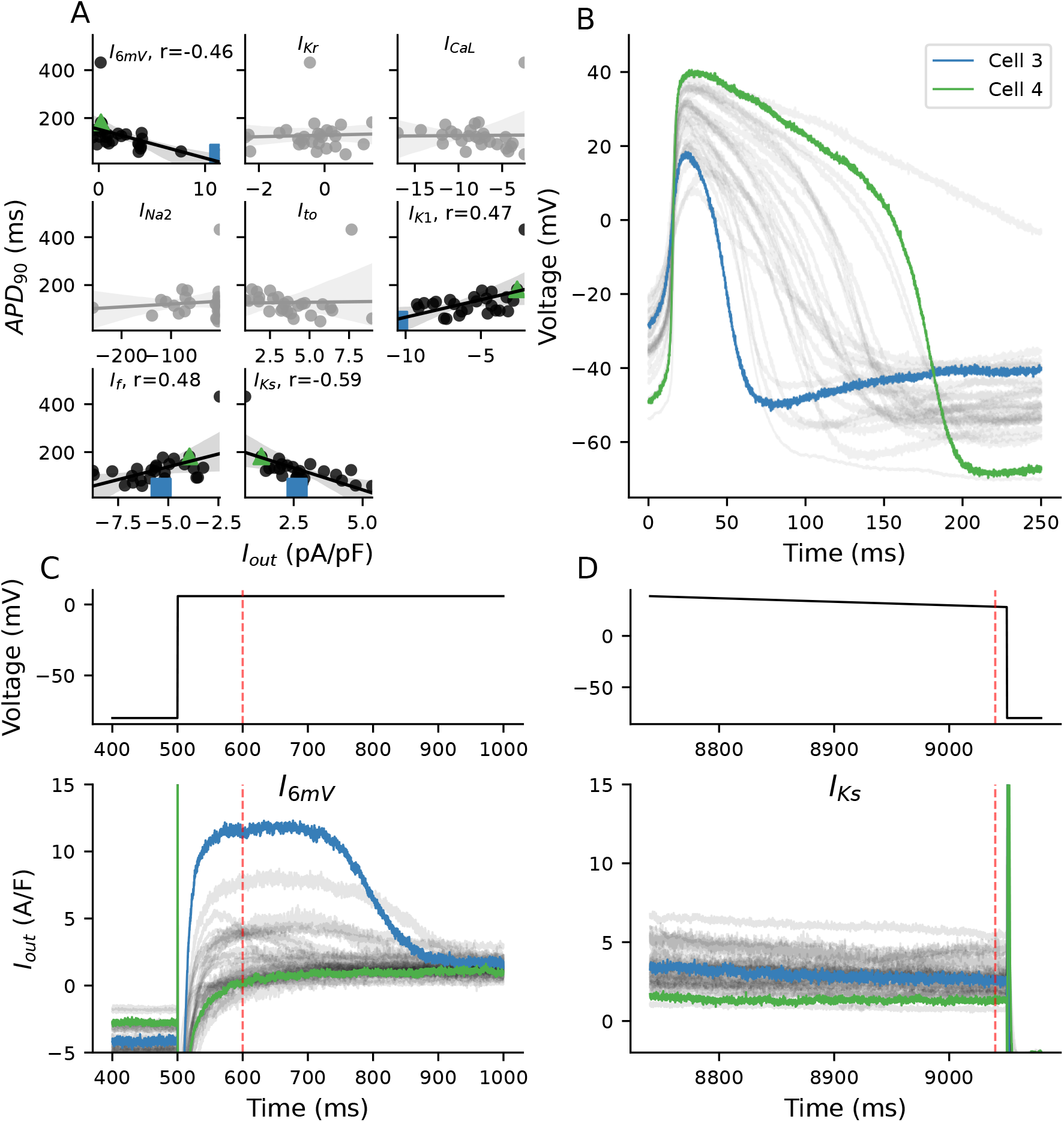
Correlations between ionic currents and APD_90_. **A**, I_out_ from the I_6mV_, I_K1_, I_f_, and I_Ks_ segments correlate significantly (p*<*.05) with APD_90_. **B**, Cells that generated the shortest (Short) and second longest (Long) APs were investigated. **C**-**D**,Current responses from cells during I_6mV_ (**C**) and I_Ks_ (**D**) step.

To illustrate the relationship between currents and APD_90_, we investigated the current responses from two different cells (Figure 10B): one with the shortest APD_90_ (Cell 3) and one with the second longest APD_90_ (Cell 4).

The strong correlation between the I_Ks_-isolating segment and APD_90_ indicates that the net outward current at large positive voltages affects APD_90_. Cell 3 has a slightly larger outward current during this I_Ks_-isolating segment when compared to Cell 4 (Figure 10D).

I_f_ and I_K1_-isolating segments correlate less strongly with APD_90_ and in the opposite direction than what would be expected. While both of these currents are expected to have an effect on AP duration, their current-isolating segments are likely contaminated with I_leak_ (Figure 9) that is contributing to the opposite direction of this (Clark et al., 2023).

The outward current present during the I_6mV_ segment is much larger in Cell 1 than any other in the population, and is very likely driving the rapid repolarization shown in this cell. In contrast, Cell 2 has a nearly balanced net current during the I_6mV_ segment. In an attempt to understand the dynamics at play, we used our population of models to study the ionic currents that may be contributing to total current during this segment.

Figure 11 shows experimental current responses (gray) and the range of traces generated from the population of models during the I_6mV_ step (600 ms). The experimental responses for many of the cells (11/39) during this I_6mV_ segment are more positive than any individual in the model population. This data indicates that several of these cells have a strong repolarizing ionic current that is not represented in the iPSC-CM electrophysiological model.

**Figure 11:**
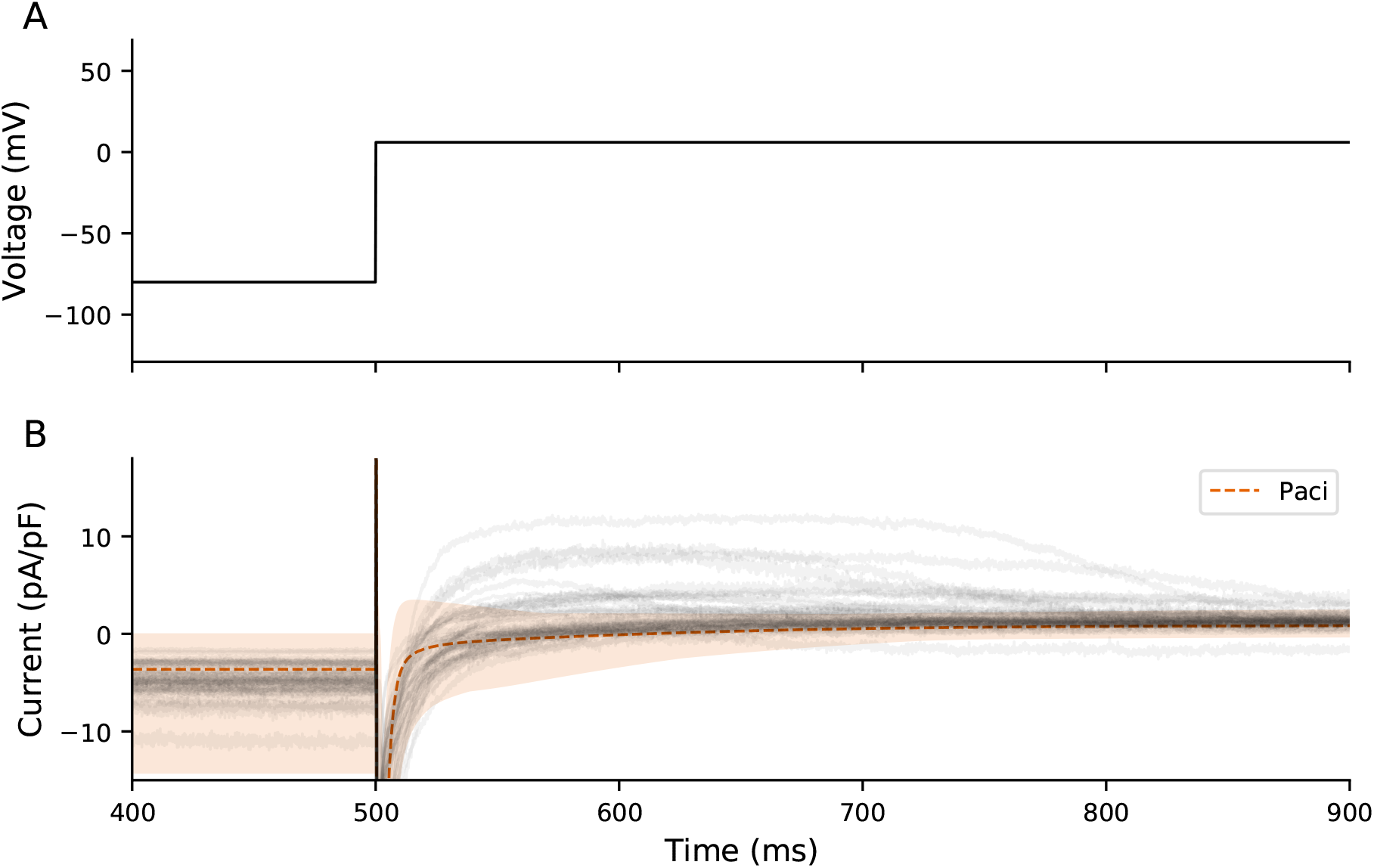
Comparison of Paci and experimental I_6mV_. Experimental (gray) and range of Paci model traces (shaded orange) during I_6mV_ voltage step.

## 4 Discussion

iPSC-CM heterogeneity is an understudied issue that confounds experimental results. Here, we propose RICP as a novel approach to study the ionic current underpinnings of AP heterogeneity. By analyzing data from a brief, optimized 10 s VC protocol, we generate the following insights about the iPSC-CMs used in this study:

- I_CaL_ is partly responsible for driving upstroke when MP is >-70 mV
- I_Kr_ contributes to establishing the MP
- I_leak_ contaminates VC responses at very positive and negative voltages and contributes to MP
- Strong outward current elicited in the range of I_Ks_ activation correlates with AP duration
- These cells have an unidentified outward current present at 6 mV that is not recapitulated by iPSC-CM mathematical models

Overall, this study demonstrates both the large cell-to-cell heterogeneity in iPSC-CMs and provides a tractable method to study the ionic current determinants of such variability during patch clamp experiments.

### 4.1 iPSC-CM heterogeneity confounds experimental results and limits reproducibility

The utility of iPSC-CMs in arrhythmia research has long been limited by their heterogeneous and unpredictable phenotype. The inherent electrophysiological variation of these cells makes it difficult to measure consistent population-level signals. The results can be confounding. For example, Blinova et al. (2019) show how iPSC-CMs purchased from the same vendor can produce opposite results to the same drug treatment protocol applied in different labs. The same group also conducted a comparative study of iPSC-CMs derived from multiple healthy individuals and showed no obvious patient-specific characteristics, likely due to the inherent heterogeneity of the cells (Blinova et al., 2019). Unfortunately, little is known about the sources or ionic current underpinnings that lead to such inconsistent results.

Shortly after the development of iPSC-CMs (Zhang et al., 2009), investigators began to sort them into nodal, atrial, and ventricular groups based on their AP features (Ma et al., 2011; Moretti et al., 2010; Itzhaki et al., 2011). The resultant sub-groups have less variance in AP morphology than the entire population of cells; i.e., sub-grouping reduces the apparent population heterogeneity. It has since been shown that, when groupings are ignored, iPSC-CM AP features are normally distributed (Du et al., 2015), leading some to argue that iPSC-CM chamber-specific grouping is a fallacy (Kane et al., 2016). While there was some initial push-back (Giles and Noble, 2016), evidence has since accumulated that indicates AP-based chamber specifications can be reductive and misleading. For example, individual iPSC-CMs do not have canonical gene expression profiles and are sensitive to noncanonical modulators of cell fate, pointing towards a physiology that is not seen in primary cardiomyocytes (Biendarra-Tiegs et al., 2019). Similarly, gene expression and phenotype data acquired from iPSC-CMs has shown a combination of attributes in a mixed population that is not consistent with any one chamber (Schmid et al., 2021).

iPSC-CM heterogeneity has long been viewed as a problem, which has led to much research into optimizing differentiation protocols to produce consistent, largely homogeneous cellular populations for *in vitro* cardiotoxicity studies (Blinova et al., 2018). We believe these characteristics of iPSC-CMs should be reframed, not as a problem that needs to be solved, but as biological reality that can serve as a laboratory model of cardiomyocyte heterogeneity.

It is well known that the AP morphology of primary cardiomyocytes isolated from the same regions of the of the heart can vary widely (Lachaud et al., 2022; Zaniboni et al., 2000). Lachaud et al. (2022) isolated 150 cells from a single rabbit left ventricle and, using optical recordings, showed that the APD_90_ ranged from 100 to >300 ms. While these primary cells specifically did not possess the multi-chamber molecular expression heterogeneity (Moretti et al., 2010) seen in iPSC-CMs, such results still indicate that substantial cell-to-cell variation in AP morphology is ubiquitous. iPSC-CMs have the potential to serve as a valuable tool to study and develop methods to help us understand electrophysiological heterogeneity with the hope of surfacing physiologically relevant patterns. We believe the work presented here serves as a step in this direction.

### 4.2 RICP is a tool to understand ionic current mechanisms of iPSC-CM heterogeneity

We used our recently published VC protocol to capture a rich snapshot of current dynamics that is explanatory of iPSC-CM AP morphology. This 10 s protocol provided information about several currents (e.g., I_CaL_, I_Kr_) and their effects on AP morphology (dV/dt_max_, MP). Other studies have collected VC data for one or two currents and shown correlations with certain AP morphological features from the same cells (Feyen et al., 2020; Schmid et al., 2020). However, we believe this is the first study that attempts to collect information about several currents with a *<*10 s protocol and use this data to explain AP features.

The VC protocol was designed to identify the presence and relative size of seven individual currents, but each of the segments is contaminated, at least to some extent, by off-target currents. For example, I_leak_ contaminates every segment of the protocol, as it contributes current at all voltages, with increased I_leak_ at voltages far from 0 mV. The sensitivity analysis shows how the I_K1_, I_to_, I_f_, and I_Ks_ segments, all of which are clamped far from 0 mV, are sensitive to changes in the seal-leak resistance (Figure 9). Some currents (e.g., I_Na_) have distinct dynamics that provide for relatively easy isolation, but others (e.g., I_Kr_) always open when several currents conduct ions.

The isolation of currents could potentially be improved through the development of new VC protocols with different optimization methods or by using new computational models that better capture the dynamics of iPSC-CMs used in a specific experimental context. However, we believe new protocols will only result in marginal improvements. Instead, we think there is far more to gain by developing methods to tease apart contributions from the different currents. Depending on the objective of the experiments, one could sequentially dissect current contributions (Banyasz et al., 2011) or fit computational models to individual traces (Groenendaal et al., 2015; Lei et al., 2017) — either of which could provide estimates of cell-specific ionic current densities.

### 4.3 I_CaL_ likely drives upstroke in many iPSC-CM studies

Using RICP we identified I_CaL_ as a contributor to upstroke velocity in the depolarized iPSC-CMs used in this study. Despite the presence of I_Na_ in most of our cells, I_CaL_ appears to be the predominant driver of upstroke. This is likely because, at depolarized MP values (e.g., > −70 mV), I_Na_ is inactive and cannot generate an inward current to drive upstroke.

This implies a mechanism similar to AP generation in SA nodal cells — despite the presence of I_Na_ (Verkerk et al., 2009), the upstroke is primarily driven by calcium currents (Vinogradova et al., 2000). This finding also hints at I_CaL_ as the likely driver of upstroke in a few of the studies in Figure 2 where dV/dt_max_ is *<* 20 V/s — e.g., the “atrial” cells from Es-Salah-Lamoureux et al. (2016). As we, and others have shown, it is possible to recover a polarized MP by dynamically clamping a synthetic I_K1_ (Goversen et al., 2018a; Becker et al., 2020; Clark et al., 2022). Such an approach makes it possible for I_Na_, among other currents, to recover from inactivation and results in APs with a faster upstroke velocity and more mature appearance.

### 4.4 I_Kr_ is likely an important current in establishing MP in depolarized iPSC-CMs

I_K1_ conductance is often thought to be reduced in iPSC-CMs relative to adult cardiomyocytes, resulting in the depolarized MP. However, iPSC-CMs can have large amounts of I_K1_, and yet, still be depolarized compared to adult cells (Horváth et al., 2018). One hypothesis for this discrepancy in MP between adult and iPSC cardiomyocytes is the increased role of I_leak_ in iPSC-CMs (Horváth et al., 2018; Clark et al., 2023) caused by their smaller size relative to adult cardiomyocytes.

In this study, we identify I_Kr_ as likely playing a role in establishing the MP of iPSC-CMs. The correlation between I_Kr_ and MP is unlikely to be substantially influenced by leak as the VC step to isolate I_Kr_ is close to the expected seal leak reversal potential. In both our experimental data and in populations of models based on either the Paci or Kernik models, the I_Kr_-isolating segment has the strongest correlation to MP (-0.66, -0.54, and -0.58 for experimental data, Paci, and Kernik, respectively, Figures 6, S3 and S4).

This finding is in agreement with previous work showing that iPSC-CM MP values are sensitive to E-4031 (Doss et al., 2012). Our modeling work indicates that with a small I_leak_ (2 GΩ seal) and substantial reduction in baseline I_Kr_, the model will depolarize to MP values similar to those shown in our study.

On a related note, I_Kr_ appears to be absent or significantly reduced when iPSC-CMs are prepared for experimentation in automated patch-clamp systems (Ismaili et al., 2023) — we hypothesize that this lack of repolarizing current may be directly responsible for the depolarized MP and lack of spontaneity in iPSC-CMs used with automated patch-clamp systems cells (Li et al., 2019).

### 4.5 The optimized VC protocol elicits a large unidentified outward current at 6 mV in some cells

One of the core features of RICP is that we use a noncanonical VC protocol to quickly probe a wide range of system dynamics. Most electrophysiological experiments, on the other hand, are designed with well-established VC protocols to study one current at a time.Such studies are effective at characterizing the magnitude and kinetics of target ionic currents. These protocols are, by design, reductive in nature. They attempt to extract information about a component part (the ion channel) from the broader dynamic system. As a result, these protocols are far less likely to identify unexpected system behaviors arising outside of that target current.

We identified a region of the protocol (stepping from -80 mV to 6 mV) that contributed information not captured in other current-isolating segments. While we are unable to identify the current present in a subset of cells during the I_6mV_ time point (Figure 10C), it appears to correlate with a shortened APD_90_. The computational models are unable to capture these dynamics, and so the mechanism remains unclear.

This could be a non-cardiac current — Horváth et al. (2020) previously identified the presence of the non-cardiac big conductance Ca^2+^-activated K^+^ current (I_BK,Ca_) in their cells. They suggest that this could have been introduced by genetic alterations (Kilpinen et al., 2017) that can occur during cell differentiation and culture. While we think this is an unlikely explanation for the current we see in our cells, it is a plausible hypothesis and provides a potential mechanism that may play a role in iPSC-CM heterogeneity that can lead to reproducibility issues.

### 4.6 Beat-to-beat variability and ionic determinants of cycle length

We did not see a significant correlation between any current-isolating element and CL (Figure S2). There was a tendency towards more current in the I_f_-isolating segment and shorter CL, as seen in the Paci-based population of models (Figure S3), but this did not meet the threshold for statistical significance (p*<*.05).

Investigation of correlations between CL and underlying ionic currents is made difficult by the substantial beat-to-beat variability of CL in some of our cells (Figure S1). The average coefficient of variation for cycle length is 13.8% while for APD_90_ it is only 3.3% and 7.1% for dV/dt_max_. For comparison, (rabbit) sinoatrial nodal cells beat much more regularly (with a coefficient of variation of 2.0% CL (Wilders and Jongsma, 1993)), as do embryonic chick ventricular cells (3.9% (Krogh-Madsen et al., 2017)). The mechanism of why some iPSC-CMs have such irregular cycle lengths with a slow and erratic diastolic phase, rather than a steady depolarization to a threshold for AP take-off, is not is not clear.

### 4.7 Limitations and future directions

The RICP method shows how a brief VC protocol can be used to provide mechanistic insights into AP morphology and heterogeneity. This protocol, however, does not perfectly isolate each of the seven currents, and so the causal relationships of currents with AP morphological features is likely weaker than if we conducted traditional drug block experiments. In the future, we believe methods that tease apart the current contributions at each time point has the potential to improve insights drawn from this VC protocol, such as ion channel co-expression patterns.

In this study, we focus on a set of cells from a single differentiation batch. While this highlighted the extent of heterogeneity within cells that should be very similar, in the future it would be interesting to conduct this RICP approach on cells from multiple donors, across multiple differentiation batches, and in multiple laboratory settings.

### 4.8 Conclusion

We believe that RICP has the potential to improve our understanding of iPSC-CM heterogeneity. The brief duration and targeting of most key cardiac ionic currents make this protocol a valuable tool in the study of iPSC-CM AP heterogeneity and drug mechanisms. If all single-cell patch-clamp iPSC-CM experiments collected this type of data, we believe the field would have a better grasp on the ionic current underpinnings of inter- and intralab heterogeneity.

## 6 Author Contributions

A.P.C. designed research, analyzed data, prepared figures, and drafted manuscript. A.P.C. and S.W. performed experiments. A.P.C. and K.E.F. completed the sensitivity analysis. A.P.C., K.E.F., T.K.M., and D.J.C. interpreted results. T.K.M. and D.J.C. edited and revised manuscript. All authors approved final version of manuscript.

## References

Akwaboah AD, Tsevi B, Yamlome P, Treat JA, Brucal-Hallare M, Cordeiro JM & Deo M (2021). An in silico hiPSC-Derived Cardiomyocyte Model Built With Genetic Algorithm. Frontiers in Physiology 12, 675867.

Banyasz T, Horvath B, Jian Z, Izu LT & Chen-Izu Y (2011). Sequential dissection of multiple ionic currents in single cardiac myocytes under action potential-clamp. Journal of Molecular and Cellular Cardiology 50, 578–581.

Becker N, Horváth A, De Boer T, Fabbri A, Grad C, Fertig N, George M & Obergrussberger A (2020). Automated Dynamic Clamp for Simulation of IK1 in Human Induced Pluripotent Stem Cell–Derived Cardiomyocytes in Real Time Using Patchliner Dynamite8. . Current Protocols in Pharmacology 88, 1–23.

Biendarra-Tiegs SM, Li X, Ye D, Brandt EB, Ackerman MJ & Nelson TJ (2019). Single-Cell RNA-Sequencing and Optical Electrophysiology of Human Induced Pluripotent Stem Cell-Derived Cardiomyocytes Reveal Discordance between Cardiac Subtype-Associated Gene Expression Patterns and Electrophysiological Phenotypes. . Stem Cells and Development 28, 659–673.

Blinova K, Dang Q, Millard D, Smith G, Pierson J, Guo L, Brock M, Lu HR, Kraushaar U, Zeng H, Shi H, Zhang X, Sawada K, Osada T, Kanda Y, Sekino Y, Pang L, Feaster TK, Kettenhofen R, Stockbridge N, Strauss DG & Gintant G (2018). International Multisite Study of Human-Induced Pluripotent Stem Cell-Derived Cardiomyocytes for Drug Proarrhythmic Potential Assessment. . Cell Reports 24, 3582–3592.

Blinova K, Schocken D, Patel D, Daluwatte C, Vicente J, Wu JC & Strauss DG (2019). Clinical Trial in a Dish: Personalized Stem Cell–Derived Cardiomyocyte Assay Compared With Clinical Trial Results for Two QT-Prolonging Drugs. . Clinical and Translational Science 12, 687–697.

Britton OJ, Bueno-Orovio A, Van Ammel K, Lu HR, Towart R, Gallacher DJ & Rodriguez B (2013). Experimentally calibrated population of models predicts and explains intersubject variability in cardiac cellular electrophysiology. . Proceedings of the National Academy of Sciences 110, E2098–E2105.

Clark AP, Clerx M, Wei S, Lei CL, de Boer TP, Mirams GR, Christini DJ & Krogh-Madsen T (2023). Leak current, even with gigaohm seals, can cause misinterpretation of stem cell-derived cardiomyocyte action potential recordings. . Europace : European pacing, arrhythmias, and cardiac electrophysiology : journal of the working groups on cardiac pacing, arrhythmias, and cardiac cellular electrophysiology of the European Society of Cardiology .

Clark AP, Wei S, Kalola D, Krogh-Madsen T & Christini DJ (2022). An in silico-in vitro pipeline for drug cardiotoxicity screening identifies ionic pro-arrhythmia mechanisms. . British journal of pharmacology 179, 4829–4843.

Clerx M, Collins P, de Lange E & Volders PGA (2016). Myokit: A simple interface to cardiac cellular electrophysiology. Progress in biophysics and molecular biology 120, 100–14.

Doss MX, Di Diego JM, Goodrow RJ, Wu Y, Cordeiro JM, Nesterenko VV, Barajas-Martínez H, Hu D, Urrutia J, Desai M, Treat JA, Sachinidis A & Antzelevitch C (2012). Maximum diastolic potential of human induced pluripotent stem cell-derived cardiomyocytes depends critically on IKr. . PLoS ONE 7.

Du DT, Hellen N, Kane C & Terracciano CM (2015). Action potential morphology of human induced pluripotent stem cell-derived cardiomyocytes does not predict cardiac chamber specificity and is dependent on cell density. . Biophysical Journal 108, 1–4.

Es-Salah-Lamoureux Z, Jouni M, Malak OA, Belbachir N, Al Sayed ZR, Gandon-Renard M, Lamirault G, Gauthier C, Baró I, Charpentier F, Zibara K, Lemarchand P, Beaumelle B, Gaborit N & Loussouarn G (2016). HIV-Tat induces a decrease in IKr and IKs via reduction in phosphatidylinositol-(4,5)-bisphosphate availability. . Journal of Molecular and Cellular Cardiology 99, 1–13.

Feyen DA, McKeithan WL, Bruyneel AA, Spiering S, Hörmann L, Ulmer B, Zhang H, Briganti F, Schweizer M, Hegyi B, Liao Z, Pölönen RP, Ginsburg KS, Lam CK, Serrano R, Wahlquist C, Kreymerman A, Vu M, Amatya PL, Behrens CS, Ranjbarvaziri S, Maas RG, Greenhaw M, Bernstein D, Wu JC, Bers DM, Eschenhagen T, Metallo CM & Mercola M (2020). Metabolic Maturation Media Improve Physiological Function of Human iPSC-Derived Cardiomyocytes. Cell Reports 32.

Giles WR & Noble D (2016). Rigorous Phenotyping of Cardiac iPSC Preparations Requires Knowledge of Their Resting Potential(s). Biophysical journal 110, 278–80.

Gong JQ & Sobie EA (2018). Population-based mechanistic modeling allows for quantitative predictions of drug responses across cell types. . npj Systems Biology and Applications 4.

Goversen B, Becker N, Stoelzle-Feix S, Obergrussberger A, Vos MA, van Veen TA, Fertig N & de Boer TP (2018a). A hybrid model for safety pharmacology on an automated patch clamp platform: Using dynamic clamp to join iPSC-derived cardiomyocytes and simulations of Ik1 ion channels in real-time. Frontiers in Physiology 8, 1–10.

Goversen B, van der Heyden MA, van Veen TA & de Boer TP (2018b). The immature electrophysiological phenotype of iPSC-CMs still hampers in vitro drug screening: Special focus on IK1. Pharmacology and Therapeutics 183, 127–136.

Grandi E, Navedo MF, Saucerman JJ, Bers DM, Chiamvimonvat N, Dixon RE, Dobrev D, Gomez AM, Harraz OF, Hegyi B, Jones DK, Krogh-Madsen T, Murfee WL, Nystoriak MA, Posnack NG, Ripplinger CM, Veeraraghavan R & Weinberg S (2023). Diversity of cells and signals in the cardiovascular system. The Journal of physiology 601, 2547–2592.

Groenendaal W, Ortega FA, Kherlopian AR, Zygmunt AC, Krogh-Madsen T & Christini DJ (2015). Cell-Specific Cardiac Electrophysiology Models. PLoS Computational Biology 11, 1–22.

Han L, Li Y, Tchao J, Kaplan AD, Lin B, Li Y, Mich-Basso J, Lis A, Hassan N, London B, Bett GC, Tobita K, Rasmusson RL & Yang L (2014). Study familial hypertrophic cardiomyopathy using patient-specific induced pluripotent stem cells. Cardiovascular Research 104, 258–269.

Horváth A, Christ T, Koivumäki JT, Prondzynski M, Zech AT, Spohn M, Saleem U, Mannhardt I, Ulmer B, Girdauskas E, Meyer C, Hansen A, Eschenhagen T & Lemoine MD (2020). Case Report on: Very Early Afterdepolarizations in HiPSC-Cardiomyocytes-An Artifact by Big Conductance Calcium Activated Potassium Current (Ibk,Ca). . Cells 9.

Horváth A, Lemoine MD, Löser A, Mannhardt I, Flenner F, Uzun AU, Neuber C, Breckwoldt K, Hansen A, Girdauskas E, Reichenspurner H, Willems S, Jost N, Wettwer E, Eschenhagen T & Christ T (2018). Low Resting Membrane Potential and Low Inward Rectifier Potassium Currents Are Not Inherent Features of hiPSC-Derived Cardiomyocytes. . Stem Cell Reports 10, 822–833.

Ismaili D, Schulz C, Horváth A, Koivumäki JT, Mika D, Hansen A, Eschenhagen T & Christ T (2023). Human induced pluripotent stem cell-derived cardiomyocytes as an electrophysiological model: Opportunities and challenges-The Hamburg perspective. . Frontiers in physiology 14, 1132165.

Itzhaki I, Maizels L, Huber I, Zwi-Dantsis L, Caspi O, Winterstern A, Feldman O, Gepstein A, Arbel G, Hammerman H, Boulos M & Gepstein L (2011). Modelling the long QT syndrome with induced pluripotent stem cells. Nature 471, 225–9.

Kane C, Du DT, Hellen N & Terracciano CM (2016). The Fallacy of Assigning Chamber Specificity to iPSC Cardiac Myocytes from Action Potential Morphology. Biophysical Journal 110, 281–283.

Kernik DC, Morotti S, Wu HD, Garg P, Duff HJ, Kurokawa J, Jalife J, Wu JC, Grandi E & Clancy CE (2019). A computational model of induced pluripotent stem-cell derived cardiomyocytes incorporating experimental variability from multiple data sources. . Journal of Physiology 597, 4533–4564.

Kilpinen H, Goncalves A, Leha A, Afzal V, Alasoo K, Ashford S, Bala S, Bensaddek D, Casale FP, Culley OJ, Danecek P, Faulconbridge A, Harrison PW, Kathuria A, McCarthy D, McCarthy SA, Meleckyte R, Memari Y, Moens N, Soares F, Mann A, Streeter I, Agu CA, Alderton A, Nelson R, Harper S, Patel M, White A, Patel SR, Clarke L, Halai R, Kirton CM, Kolb-Kokocinski A, Beales P, Birney E, Danovi D, Lamond AI, Ouwehand WH, Vallier L, Watt FM, Durbin R, Stegle O & Gaffney DJ (2017). Common genetic variation drives molecular heterogeneity in human iPSCs. Nature 546, 370–375.

Krogh-Madsen T, Kold Taylor L, Skriver AD, Schaffer P & Guevara MR (2017). Regularity of beating of small clusters of embryonic chick ventricular heart-cells: experiment vs. stochastic single-channel population model. . Chaos (Woodbury, N.Y.) 27, 093929.

Krogh-Madsen T, Sobie EA & Christini DJ (2016). Improving cardiomyocyte model fidelity and utility via dynamic electrophysiology protocols and optimization algorithms. . Journal of Physiology 594, 2525–2536.

Lachaud Q, Aziz MHN, Burton FL, Macquaide N, Myles RC, Simitev RD & Smith GL (2022). Electrophysiological heterogeneity in large populations of rabbit ventricular cardiomyocytes. Cardiovascular Research pp. 3112–3125.

Lee JH, Protze SI, Laksman Z, Backx PH & Keller GM (2017). Human Pluripotent Stem Cell-Derived Atrial and Ventricular Cardiomyocytes Develop from Distinct Mesoderm Populations. . Cell Stem Cell 21, 179–194.

Lei CL (2020). Model-Driven Design and Uncertainty Quantification for Cardiac Electrophysiology Experiments Ph.D. diss., University of Oxford.

Lei CL, Clerx M, Whittaker DG, Gavaghan DJ, de Boer TP & Mirams GR (2020). Accounting for variability in ion current recordings using a mathematical model of artefacts in voltage-clamp experiments. . Philosophical transactions. Series A, Mathematical, physical, and engineering sciences 378, 20190348.

Lei CL, Wang K, Clerx M, Johnstone RH, Hortigon-Vinagre MP, Zamora V, Allan A, Smith GL, Gavaghan DJ, Mirams GR & Polonchuk L (2017). Tailoring mathematical models to stem-cell derived cardiomyocyte lines can improve predictions of drug-induced changes to their electrophysiology. . Frontiers in Physiology 8, 1–13.

Li W, Luo X, Ulbricht Y, Wagner M, Piorkowski C, El-Armouche A & Guan K (2019). Establishment of an automated patch-clamp platform for electrophysiological and pharmacological evaluation of hiPSC-CMs. . Stem Cell Research 41, 101662.

Liang P, Sallam K, Wu H, Li Y, Itzhaki I, Garg P, Zhang Y, Vermglinchan V, Lan F, Gu M, Gong T, Zhuge Y, He C, Ebert AD, Sanchez-Freire V, Churko J, Hu S, Sharma A, Lam CK, Scheinman MM, Bers DM & Wu JC (2016). Patient-Specific and Genome-Edited Induced Pluripotent Stem Cell-Derived Cardiomyocytes Elucidate Single-Cell Phenotype of Brugada Syndrome. . Journal of the American College of Cardiology 68, 2086–2096.

Ma J, Guo L, Fiene SJ, Anson BD, Thomson JA, Kamp TJ, Kolaja KL, Swanson BJ, January CT, Kl K, Bj S & Ct J (2011). High purity human-induced pluripotent stem cell-derived cardiomyocytes : electrophysiological properties of action potentials and ionic currents. . Am J Physiol Heart Circ Physiol 301, 2006–2017.

Mathur A, Loskill P, Shao K, Huebsch N, Hong SG, Marcus SG, Marks N, Mandegar M, Conklin BR, Lee LP & Healy KE (2015). Human iPSC-based cardiac microphysiological system for drug screening applications. Scientific Reports 5, 1–7.

Moretti A, Bellin M, Welling A, Jung CB, Lam JT, Bott-Flügel L, Dorn T, Goedel A, Höhnke C, Hofmann F, Seyfarth M, Sinnecker D, Schömig A & Laugwitz KL (2010). Patient-specific induced pluripotent stem-cell models for long-QT syndrome. The New England journal of medicine 363, 1397–409.

Ni H, Morotti S & Grandi E (2018). A Heart for Diversity: Simulating Variability in Cardiac Arrhythmia Research. Frontiers in Physiology 9, 958.

Paci M, Hyttinen J, Aalto-Setälä K & Severi S (2013). Computational Models of Ventricular- and Atrial-Like Human Induced Pluripotent Stem Cell Derived Cardiomyocytes. . Annals of Biomedical Engineering 41, 2334–2348.

Passini E, Britton OJ, Lu HR, Rohrbacher J, Hermans AN, Gallacher DJ, Greig RJ, Bueno-Orovio A & Rodriguez B (2017). Human in silico drug trials demonstrate higher accuracy than animal models in predicting clinical pro-arrhythmic cardiotoxicity. . Frontiers in Physiology 8, 1–15.

Quach B, Krogh-Madsen T, Entcheva E & Christini DJ (2018). Light-Activated Dynamic Clamp Using iPSC-Derived Cardiomyocytes. Biophysical Journal 115, 2206–2217.

Rook MB, Alshinawi CB, Groenewegen WA, Van Gelder IC, Van Ginneken AC, Jongsma HJ, Mannens MM & Wilde AA (1999). Human SCN5A gene mutations alter cardiac sodium channel kinetics and are associated with the Brugada syndrome. . Cardiovascular Research 44, 507–517.

Sarkar AX, Christini DJ & Sobie EA (2012). Exploiting mathematical models to illuminate electrophysiological variability between individuals. Journal of Physiology 590, 2555–2567.

Schmid C, Abi-Gerges N, Leitner MG, Zellner D & Rast G (2021). Ion channel expression and electrophysiology of singular human (Primary and induced pluripotent stem cell-derived) cardiomyocytes. . Cells 10.

Schmid C, Wohnhaas CT, Hildebrandt T, Baum P & Rast G (2020). Characterization of iCell cardiomyocytes using single-cell RNA-sequencing methods. Journal of pharmacological and toxicological methods 106, 106915.

Terrenoire C, Wang K, Chan Tung KW, Chung WK, Pass RH, Lu JT, Jean JC, Omari A, Sampson KJ, Kotton DN, Keller G & Kass RS (2013). Induced pluripotent stem cells used to reveal drug actions in a long QT syndrome family with complex genetics. . Journal of General Physiology 141, 61–72.

Verkerk AO, Wilders R, Van Borren MM & Tan HL (2009). Is sodium current present in human sinoatrial node cells? International Journal of Biological Sciences 5, 201–204.

Vinogradova TM, Zhou YY, Bogdanov KY, Yang D, Kuschel M, Cheng H & Xiao RP (2000). Sinoatrial node pacemaker activity requires Ca(2+)/calmodulin-dependent protein kinase II activation. . Circulation research 87, 760–7.

Virtanen P, Gommers R, Oliphant TE, Haberland M, Reddy T, Cournapeau D, Burovski E, Peterson P, Weckesser W, Bright J, van der Walt SJ, Brett M, Wilson J, Millman KJ, Mayorov N, Nelson ARJ, Jones E, Kern R, Larson E, Carey CJ, Polat I, Feng Y, Moore EW, VanderPlas J, Laxalde D, Perktold J, Cimrman R, Henriksen I, Quintero EA, Harris CR, Archibald AM, Ribeiro AH, Pedregosa F, van Mulbregt P & SciPy 1.0 Contributors (2020). {SciPy} 1.0: Fundamental Algorithms for Scientific Computing in Python. Nature Methods 17, 261–272.

Whittaker DG, Clerx M, Lei CL, Christini DJ & Mirams GR (2020). Calibration of ionic and cellular cardiac electrophysiology models. Wiley interdisciplinary reviews. Systems biology and medicine 12, e1482.

Wilders R & Jongsma HJ (1993). Beating irregularity of single pacemaker cells isolated from the rabbit sinoatrial node. Biophysical journal 65, 2601–13.

Zaniboni M, Pollard AE, Yang L & Spitzer KW (2000). Beat-to-beat repolarization variability in ventricular myocytes and its suppression by electrical coupling. . American Journal of Physiology -Heart and Circulatory Physiology 278, 677–687.

Zhang J, Wilson GF, Soerens AG, Koonce CH, Yu J, Palecek SP, Thomson JA & Kamp TJ (2009). Functional cardiomyocytes derived from human induced pluripotent stem cells. Circulation research 104, 30–41.

