## Supplemental Figures for "Rapid ionic current phenotyping (RICP) identifies mechanistic underpinnings of iPSC-CM AP heterogeneity"

---

### 1 Supplementary figures

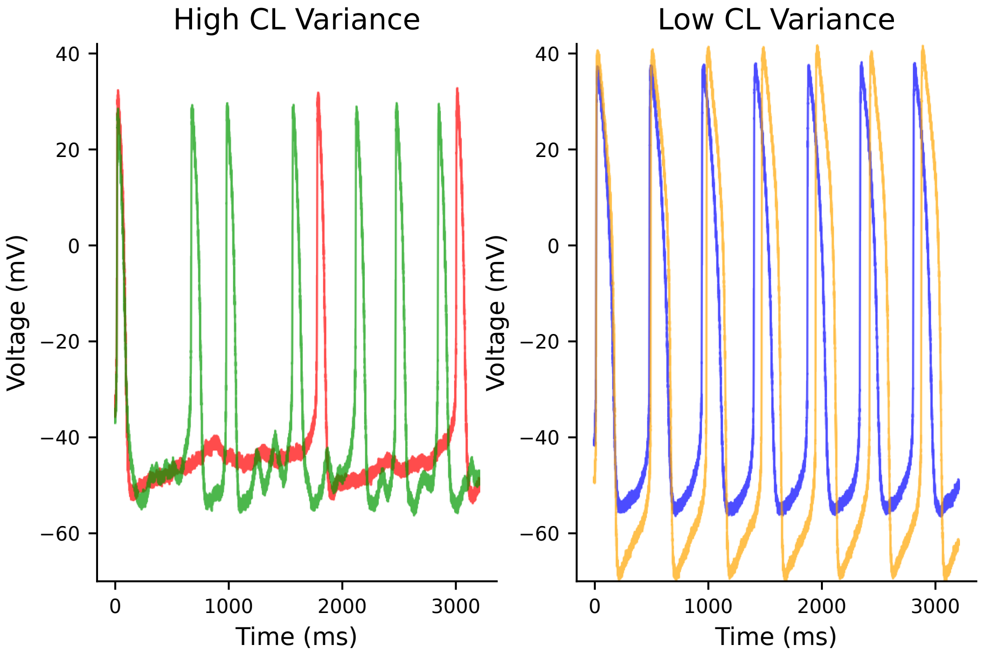


Figure 1: **Cells with high (left) and low (right) variance in cycle length**. High-variance cells have an irregular diastolic depolarization ramp and varying takeoff potential, while low-variance cells show consistent, very regular AP morphologies.

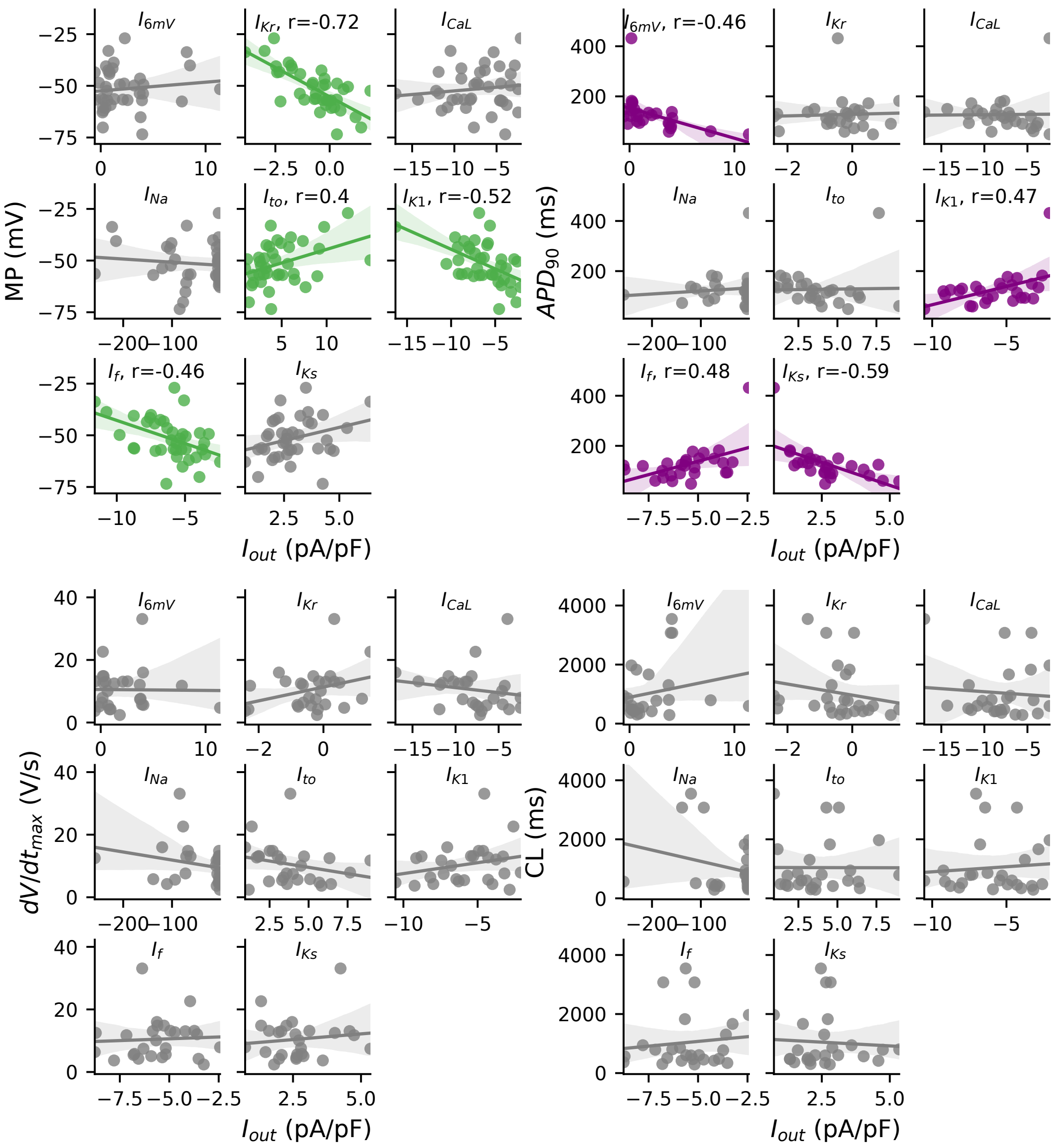


Figure 2: **AP parameter and current correlations for *in vitro* cells**. Correlations between AP metrics and current-isolating segments. Non-gray colors indicate correlations that met threshold for statistical significance.

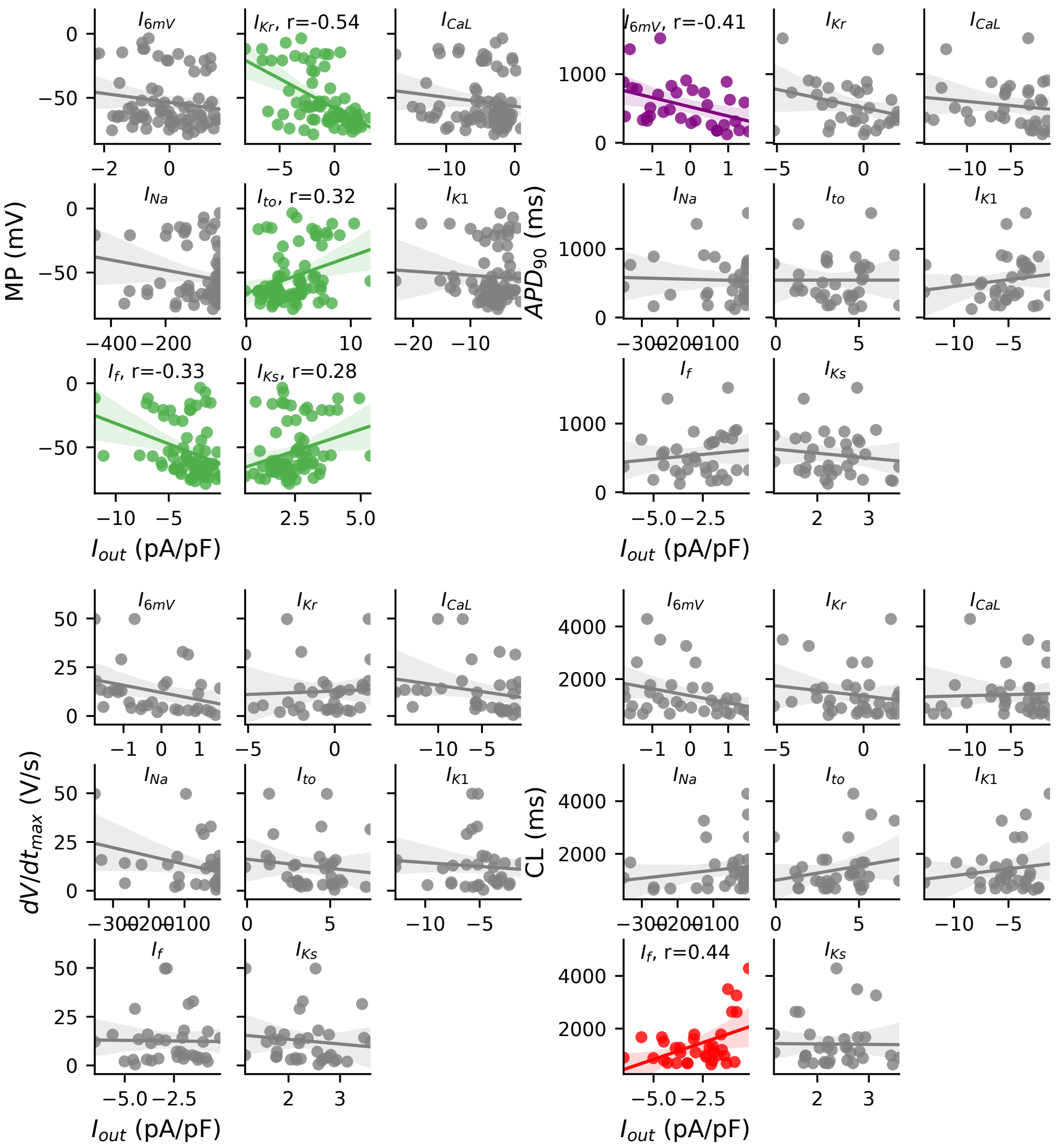


Figure 3: **AP parameter and current correlations for Paci-based model population**. Correlations between AP metrics and current-isolating segments. Non-gray colors indicate correlations that met threshold for statistical significance. There are no statistically significant correlations between $dV/dt_{max}$ and any of the current-isolating segments. $I_{Kr}$ is the main determinant of MDP, $I_f$ the main determinant of CL, and $I_{6mV}$ the main determinant of $APD_{90}$.

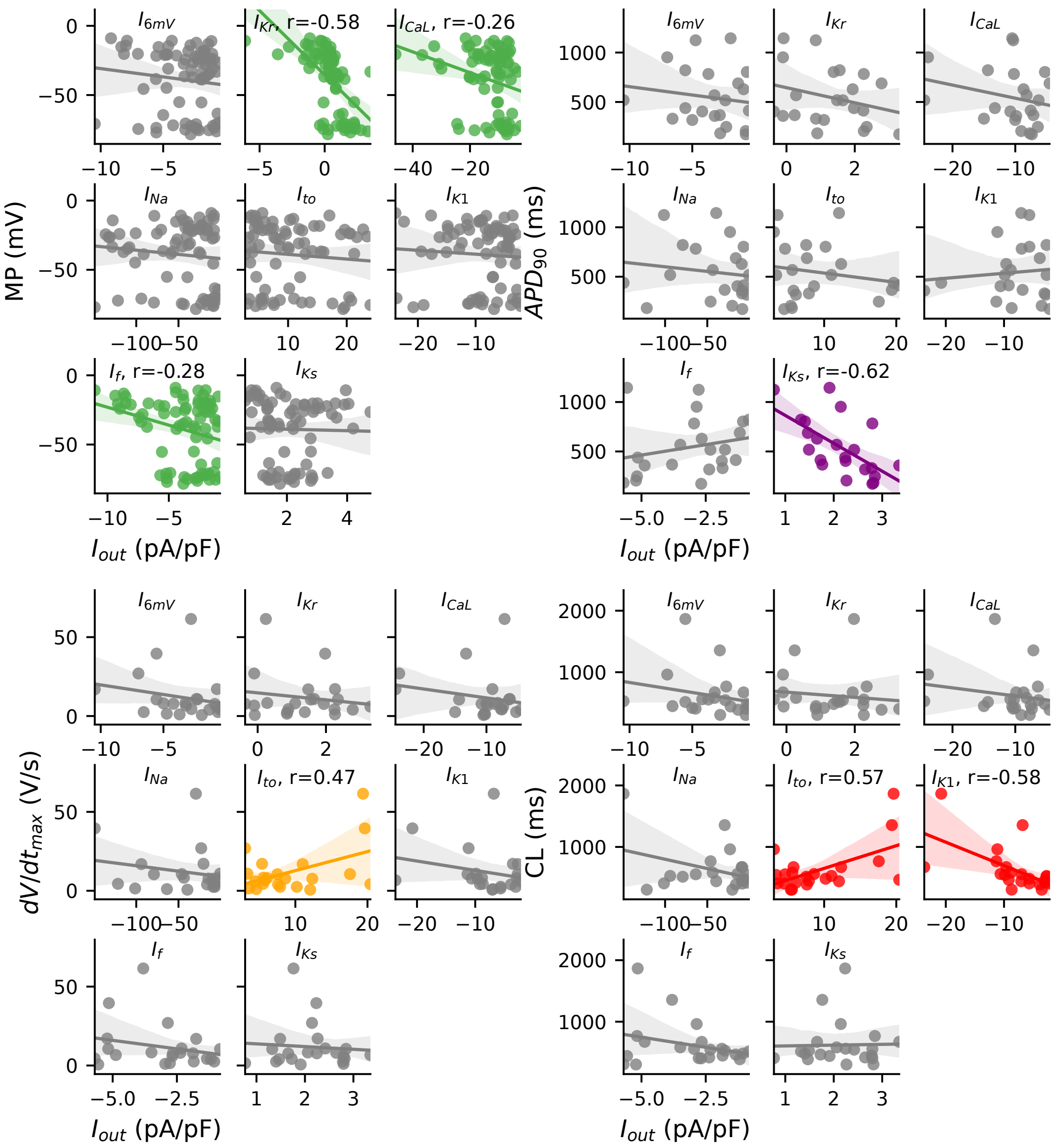


Figure 4: **AP parameter and current correlations for Kernik-based model population**. Correlations between AP metrics and current-isolating segments. Non-gray colors indicate correlations that met threshold for statistical significance. $I_{to}$ correlate with $dV/dt_{max}$, $I_{Ks}$ with $APD_{90}$, and both $I_{to}$ and $I_{K1}$ with CL. $I_{Kr}$ is the main determinant of MDP.
